# Patient-specific network connectivity combined with a next generation neural mass model to test clinical hypothesis of seizure propagation

**DOI:** 10.1101/2021.01.15.426839

**Authors:** Moritz Gerster, Halgurd Taher, Antonín Škoch, Jaroslav Hlinka, Maxime Guye, Fabrice Bartolomei, Viktor Jirsa, Anna Zakharova, Simona Olmi

## Abstract

Dynamics underlying epileptic seizures span multiple scales in space and time, therefore, understanding seizure mechanisms requires identifying the relations between seizure components within and across these scales, together with the analysis of their dynamical repertoire. In this view, mathematical models have been developed, ranging from single neuron to neural population.

In this study we consider a neural mass model able to exactly reproduce the dynamics of heterogeneous spiking neural networks. We combine the mathematical modelling with structural information from non-invasive brain imaging, thus building large-scale brain network models to explore emergent dynamics and test clinical hypothesis. We provide a comprehensive study on the effect of external drives on neuronal networks exhibiting multistability, in order to investigate the role played by the neuroanatomical connectivity matrices in shaping the emergent dynamics. In particular we systematically investigate the conditions under which the network displays a transition from a low activity regime to a high activity state, which we identify with a seizure-like event. This approach allows us to study the biophysical parameters and variables leading to multiple recruitment events at the network level. We further exploit topological network measures in order to explain the differences and the analogies among the subjects and their brain regions, in showing recruitment events at different parameter values.

We demonstrate, along the example of diffusion-weighted magnetic resonance imaging (MRI) connectomes of 20 healthy subjects and 15 epileptic patients, that individual variations in structural connectivity, when linked with mathematical dynamic models, have the capacity to explain changes in spatiotemporal organization of brain dynamics, as observed in network-based brain disorders. In particular, for epileptic patients, by means of the integration of the clinical hypotheses on the epileptogenic zone (EZ), i.e. the local network where highly synchronous seizures originate, we have identified the sequence of recruitment events and discussed their links with the topological properties of the specific connectomes. The predictions made on the basis of the implemented set of exact mean-field equations turn out to be in line with the clinical pre-surgical evaluation on recruited secondary networks.

## 1 INTRODUCTION

Epilepsy is a chronic neurological disorder characterized by the occurrence and recurrence of seizures and represents the third most common neurological disorder affecting more than 50 million people worldwide. Anti-epileptic drugs are the first line of treatment for epilepsy and they provide sufficient seizure control in around two-thirds of cases Kwan and Brodie (2000). However, about 30 to 40% of epilepsy patients do not respond to drugs, a percentage that has remained relatively stable despite significant efforts to develop new anti-epileptic medication over the past decades. For drug-resistent patients, a possible treatment is the surgical resection of the brain tissue responsible for the generation of seizures.

As a standard procedure, epilepsy surgery is preceded by a qualitative assessment of different brain imaging modalities in order to identify the brain tissue responsible for seizure generation, i.e. the epileptogenic zone (EZ) Rosenow and Lüders (2001), which in general represents a localized region or network where seizures arise, before recruiting secondary networks, called the propagation zone (PZ) Talairach and Bancaud (1966); Bartolomei et al. (2001); Spencer (2002). Outcomes are positive whenever the patient has become seizure-free after surgical operation.

Intracranial electroencephalography (iEEG) is commonly used during the presurgical assessment to find the seizure onset zone David et al. (2011); Duncan et al. (2016); Rosenow and Lüders (2001), the assumption being that the region where seizures emerge, is at least part of the brain tissue responsible for seizure generation. As a part of the standard presurgical evaluation with iEEG, stereotactic EEG (SEEG) is used to help correctly delineating the EZ Bartolomei et al. (2002). Alternative imaging techniques such as structural MRI, M/EEG, and positron emission tomography (PET) help the clinician to outline the EZ. Recently, diffusion MRI (dMRI) started being evaluated as well, thus giving the possibility to infer the connectivity between different brain regions and revealing reduced fractional anisotropy Ahmadi et al. (2009); Bernhardt et al. (2013) and structural alterations in the connectome of epileptic patients Bonilha et al. (2012); DeSalvo et al. (2014); Besson et al. (2014). However, epilepsy surgery is often unsuccessful and the long-term positive outcome may be lower than 25% in extra-temporal cases De Tisi et al. (2011); Najm et al. (2013), thus meaning that the EZ has not been correctly identified or that the EZ and the seizure onset zone may not coincide.

In order to quantitatively examine clinical data and to determine targets for surgery, many computational models have been recently proposed Hutchings et al. (2015); Goodfellow et al. (2017); Khambhati et al. (2016); Lopes et al. (2017); Sinha et al. (2017), that use MRI or iEEG data acquired during presurgical workup to infer structural or functional brain networks. Taking advantages of recent advances in our understanding of epilepsy, that indicate that seizures may arise from distributed ictogenic networks Richardson (2012); Bartolomei et al. (2017); Besson et al. (2017), phenomenological models of seizure transitions are used to compute the escape time, i.e., the time that each network node takes to transit from a normal state to a seizure-like state. Nodes with the lowest escape time are then considered as representative of the seizure onset zone and therefore candidates for surgical resection, by assuming seizure onset zone as a proxy for the EZ Hutchings et al. (2015); Sinha et al. (2017). Alternatively, different possible surgeries are simulated in silico to predict surgical outcomes Goodfellow et al. (2017); Lopes et al. (2017, 2019) by making use of synthetic networks and phenomenological network models of seizure generation. Further attention has been paid to studying how network structure and tissue heterogeneities underpin the emergence of focal and widespread seizure dynamics in synthetic networks of phase oscillators Lopes et al. (2019, 2020).

More in general there is a vast and valuable literature on computational modeling in epilepsy, where two classes of models are used: 1) mean-field (macroscopic) models and 2) detailed (microscopic) network models. Mean field models are often preferred over the more detailed models since they have fewer parameters and thus simplify the study of transitions from interictal to ictal states and the subsequent EEG analysis of data from epilepsy patients. This is justified as the macroelectrodes used for EEG recordings represent the average local field potential arising from neuronal populations. Indeed, much effort has been made so far to explain the biophysical and dynamical nature of seizure onsets and offsets by employing neural mass models Da Silva et al. (1974); Wendling et al. (2002); Kalitzin et al. (2010); Touboul et al. (2011); Kramer et al. (2012); Jirsa et al. (2014). Mechanistic interpretability of the mean field parameters is lost, as many physiological details are absorbed in few degrees of freedom. Since the mean field models remain relatively simple, they can also be employed to describe epileptic processes occurring in “large-scale” systems, e.g. the precise identification of brain structures that belong to the seizure-triggering zone (epileptic activity often spreads over quite extended regions and involves several cortical and sub-cortical structures). However, only recently, propagation of epileptic seizures started to be studied using brain network models, and was limited to small scales Terry et al. (2012) or absence seizures Taylor et al. (2013), while partial seizures have been reported to propagate through large-scale networks in humans Bartolomei et al. (2013) and animal models Toyoda et al. (2013). All in all, even though neural mass models are in general easier to analyze numerically because relatively few variables and parameters are involved, they drastically fail to suggest molecular and cellular mechanisms of epileptogenesis.

On the other hand, detailed network models are best suited for understanding the molecular and cellular bases of epilepsy and thus they may be used to suggest therapeutics targeting molecular pathways Destexhe and Sejnowski (1995); Van Drongelen et al. (2005); Turrigiano (2008); Cressman et al. (2009); Ullah et al. (2009). Due to the substantial complexity of neuronal structures, relatively few variables and parameters can be accessed at any time experimentally. Although biophysically explicit modeling is the primary technique to look into the role played by experimentally inaccessible variables in epilepsy, the usefulness of detailed biophysical models is limited by constraints in computational power, uncertainties in detailed knowledge of neuronal systems, and the required simplification for the numerical analysis. Therefore an intermediate “across-scale” approach, establishing relationships between sub-cellular/cellular variables of detailed models and mean-field parameters governing macroscopic models, might be a promising strategy to cover the gaps between these two modeling approaches Brocke et al. (2016); Schirner et al. (2018); Lindroos et al. (2018).

In view of developing a cross-scale approach, it is important to point out that large-scale brain network models emphasize the network character of the brain and merge structural information of individual brains with mathematical modeling, thus constituting in-silico approaches for the exploration of causal mechanisms of brain function and clinical hypothesis testing Proix et al. (2017, 2018); Olmi et al. (2019). In particular, in brain network models, a network region is a neural mass model of neural activity, connected to other regions via a connectivity matrix representing fiber tracts of the human brain. This form of virtual brain modeling Fuchs et al. (2000); Jirsa et al. (2002, 2010) exploits the explanatory power of network connectivity imposed as a constraint upon network dynamics and has provided important insights into the mechanisms underlying the emergence of asynchronous and synchronized dynamics of wakefulness and slow-wave sleep Goldman et al. (2020) while revealing the whole-brain mutual coupling between the neuronal and the neurotransmission systems to understand the flexibility of human brain function despite having to rely on fixed anatomical connectivity Kringelbach et al. (2020). Recent studies have pointed out the influence of individual structural variations of the connectome upon the large-scale brain network dynamics of the models, by systematically testing the virtual brain approach along the example of epilepsy Proix et al. (2017, 2018); Olmi et al. (2019). The employment of patient-specific virtual brain models derived from diffusion MRI may have a substantial impact for personalized medicine, allowing for an increase in predictivity concerning the pathophysiology of brain disorders, and their associated abnormal brain imaging patterns. More specifically a personalized brain network model derived from non-invasive structural imaging data would allow for testing of clinical hypotheses and exploration of novel therapeutic approaches.

In order to exploit the predictive power of personalized brain network models, we have implemented, on subject-specific connectomes, a next generation neural mass model that, differently from the previous applied heuristic mean-field models Proix et al. (2017, 2018); Olmi et al. (2019), is exactly derived from an infinite size network of quadratic integrate-and-fire neurons Montbrió et al. (2015), that represent the normal form of Hodgkin’s class I excitable membranes Ermentrout and Kopell (1986). This next generation neural mass model is able to describe the variation of synchrony within a neuronal population, which is believed to underlie the decrease or increase of power seen in given EEG frequency bands while allowing for a more direct comparison with the results of electrophysiological experiments like local field potential, EEG and event-related potentials (ERPs), thanks to its ability to capture the macroscopic evolution of the mean membrane potential. Most importantly, the exact reduction dimension techniques at the basis of the next generation neural mass model have been developed for coupled phase oscillators Ott and Antonsen (2008) and allow for an exact (analytical) moving upwards through the scales: While keeping the influence of smaller scales on larger ones they level out their inherent complexity. In this way it is therefore possible to develop an intermediate “across-scale” approach exploiting the 1:1 correspondence between microscopic and mesoscopic level that allows for a more detailed modelling parameters and for mapping the microscopic results to the relative ones in the regional mean field parameters.

The next generation neural mass model developed by Montbrió et al. (2015), has been recently extended to take into account time-delayed synaptic coupling Pazó and Montbrió (2016); Devalle et al. (2018) and, when integrated in a large-scale brain network, time delays in the interaction between the different brain areas, due to the finite conduction speed along fiber tracts of different lengths Rabuffo et al. (2020). The time delay, together with the effective stochasticity of the investigated dynamics give rise, both on structural connectivity matrices of mice and healthy subjects, to preferred spatiotemporal pattern formation Jirsa (2008); Petkoski and Jirsa (2020) and short-lived neuronal cascades that form spontaneously and propagate through the network under conditions of near-criticality Rabuffo et al. (2020). The largest neuronal cascades produce short-lived but robust co-fluctuations at pairs of regions across the brain, thus contributing to the organization of the slowly evolving spontaneous fluctuations in brain dynamics at rest. The introduction of extrinsic or endogenous noise sources in the framework of exact neural mass models is possible in terms of (pseudo)-cumulants expansion in Tyulkina et al. (2018); Goldobin et al. (2021).

In this paper, we have built brain network models for a cohort of 20 healthy subjects and 15 epileptic patients, implementing for each brain region a neural mass model developed by Montbrió et al. (2015). As paradigms for testing the spatiotemporal organization, we have systematically simulated the individual seizure-like propagation patterns, looking for the role played by the individual structural topologies in determining the recruitment mechanisms. Specific attention has been devoted to the analogies and differences among the self-emergent dynamics in healthy and epilepsy-affected subjects. Furthermore, for epileptic patients, we have validated the model against the presurgical stereotactic electroencephalography (SEEG) data and the standard-of-care clinical evaluation. More specifically Sec. 2 is devoted to the description of the implemented model and the applied methods. In Sec. 3.1 are reported the results specific for healthy subjects, while in Sec. 3.2 is reported a detailed analysis performed on epileptic patients. Finally a discussion on the presented results is reported in Sec. 4.

## 2 METHODS

### 2.1 Network Model

The membrane potential dynamics of the *i*-th QIF neuron in a network of size *N* can be written as

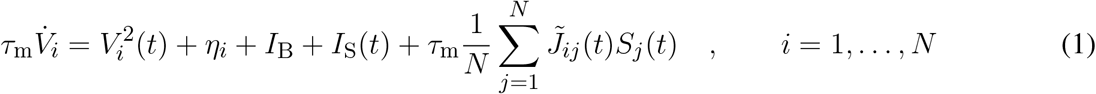

where *τ*_m_ = 20 ms is the membrane time constant and 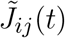 the strength of the direct synapse from neuron *j* to *i* that we assume to be constant and all identical, i.e. 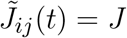. The sign of *J* determines if the neuron is excitatory (*J >* 0) or inhibitory (*J <* 0); in the following we will consider only excitatory neurons. Moreover, *η_i_* represents the neuronal excitability, *I*_B_ a constant background DC current (in the following we assume *I*_B_ = 0), *I*_S_(*t*) an external stimulus and the last term on the right hand side the synaptic current due to the recurrent connections with the pre-synaptic neurons. For instantaneous post-synaptic potentials (corresponding to *δ*-spikes) the neural activity *S_j_*(*t*) of neuron *j* reads as

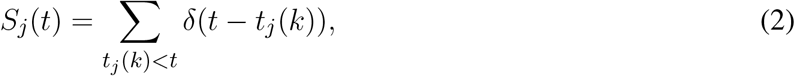

where *S_j_*(*t*) is the spike train produced by the *j*-th neuron and *t_j_*(*k*) denotes the *k*-th spike time in such sequence. We have considered a fully coupled network without autapses, therefore the post-synaptic current will be the same for each neuron.

In the absence of synaptic input, external stimuli and *I*_B_ = 0, the QIF neuron exhibits two possible dynamics, depending on the sign of *η_i_*. For negative *η_i_*, the neuron is excitable and for any initial condition 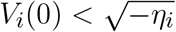, it reaches asymptotically the resting value 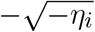. On the other hand, for initial values larger than the excitability threshold, 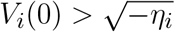, the membrane potential grows unbounded and a reset mechanism has to be introduced to describe the spiking behaviour of a neuron. Whenever *V_i_*(*t*) reaches a threshold value *V*_p_, the neuron *i* delivers a spike and its membrane potential is reset to *V*_r_, for the QIF neuron *V*_p_ = −*V*_r_ = ∞. For positive *η_i_* the neuron is supra-threshold and it delivers a regular train of spikes with frequency 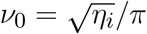.

### 2.2 Neural Mass Model

For the heterogeneous QIF network with instantaneous synapses (Eqs. (1)-(2)), an exact neural mass model has been derived in Montbrió et al. (2015). The analytic derivation is possible for QIF spiking networks using the Ott-Antonsen Ansatz Ott and Antonsen (2008) applicable to phase-oscillators networks, whenever the natural frequencies are distributed according to a Lorentzian distribution. In the case of the QIF network this corresponds to a distribution of the excitabilities {*η_i_*} given by

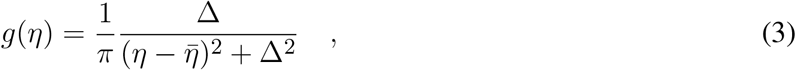

which is centred in 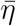 and has half width at half maximum (HWHM) Δ (Δ = 1 throughout this work). In particular, this neural mass model allows for an exact macroscopic description of the population dynamics, in the thermodynamic limit *N* → ∞, in terms of only two collective variables, namely the mean membrane voltage potential *v*(*t*) and the instantaneous population rate *r*(*t*), as follows

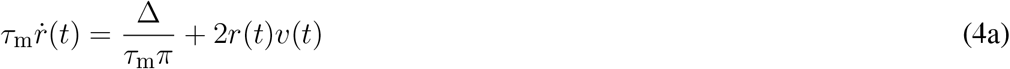

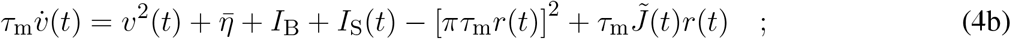

where the synaptic strength is assumed to be identical for all neurons and for instantaneous synapses in absence of plasticity 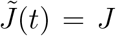. However, by including a dynamical evolution for the synapses and therefore additional collective variables, this neural mass model can be extended to any generic post-synaptic potential, see e.g. Devalle et al. (2017) for exponential synapses or Coombes and Byrne (2019) for conductance based synapses with *α*-function profile.

### 2.3 Multipopulation Neural Mass Model

The neural mass model can be easily extended to account for multiple interconnected neuronal populations *N*_pop_. In the following we consider personalized brain models derived from structural data of magnetic resonance imaging (MRI) and Diffusion Tensor weighted Imaging (DTI), thus implementing different structural connectivity matrices for healthy subjects and epileptic patients. For healthy subjects cortical and volumetric parcellations were performed using the Automatic Anatomical Atlas 1 (AAL1) (Tzourio-Mazoyer et al., 2002) with *N*_pop_ = 90 brain regions: each region will be described in terms of the presented neural mass model. For epileptic subjects cortical and volumetric parcellations were performed using the Desikan-Killiany atlas with 70 cortical regions and 17 subcortical regions (Desikan et al., 2006) (one more empty region is added in the construction of the structural connectivity for symmetry). In this case the structural connectivity matrix is composed, for each epileptic patient, by 88 nodes equipped with the presented region specific neural mass model capable of demonstrating epileptiform discharges.

The corresponding multi-population neural mass model can be straightforwardly written as

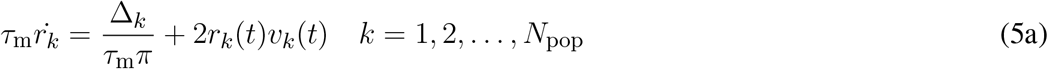

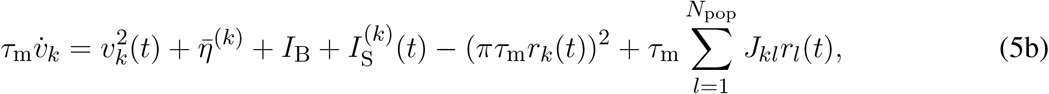

where {*J_kl_*} is the connectivity matrix, representing the synaptic weights among the populations. Diagonal entries *J_kk_* denote intra-population and non-diagonal entries *J_kl_, k* ≠ *l* inter-population connections. Here we have assumed that the neurons are globally coupled both at the intra- and inter-population level, hence removing the dependency on the neuron indices.

The connectivity matrix entries *J_kl_* are determined via a second matrix 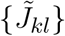, which represents the topology extracted from empirical DTI. The values of 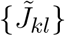 are normalized in the range [0, 1] via rescaling with the maximal entry value, and have 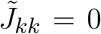 on the diagonal. In order to account for strong intra-coupling (recurrent synapses) and intermediate inter-coupling, we choose the entries of each structural connectivity as

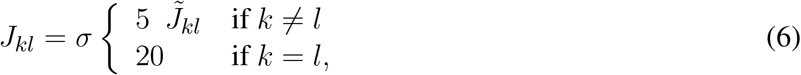

where *σ* is a rescaling factor common to all synapses, that we assume to be constant and equal to 1, apart few cases where we investigate the dependence on the synaptic weights. Hence, the synaptic weights for *k* ≠ *l* are in the range *J_kl_* ∈ [0, 5], while the intra-coupling is set to *J_kk_* = 20 (apart when specified otherwise). The time dependent stimulus current 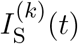 is population specific and a single population at a time is generally stimulated during a numerical experiment. The applied stimulus 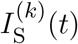 consists of a rectangular pulse of amplitude *I_S_* and duration *t_I_*; the dependence on these parameters is studied in this paper to support the generality of the results.

### 2.4 Topologies

As a first set of data, we have selected 20 diffusion-weighted magnetic resonance imaging connectomes of healthy subjects (mean age 33 years, standard deviation 5.7 years, 10 females, 2 left-handed) that participated in a study on schizophrenia as a control group (Melicher et al., 2015). All subjects were recruited via local advertisements and had none of the following conditions: Personal lifetime history of any psychiatric disorder or substance abuse established by the Mini-International Neuropsychiatric Interview (M.I.N.I.) (Lecrubier et al., 1997), any psychotic disorder in first or second-degree relatives. Further exclusion criteria included current neurological disorders, lifetime history of seizures or head injury with altered consciousness, intracranial hemorrhage, neurological sequelae, history of mental retardation, history of substance dependence, any contraindication for MRI scanning.

The scans were performed on a 3T Siemens scanner in the Institute of Clinical and Experimental Medicine in Prague, employing a Spin-Echo EPI sequence with 30 diffusion gradient directions, *TR* = 8300 ms, *TE* = 84 ms, 2 × 2 × 2*mm*^3^ voxel size, b-value 900*s/mm*^2^. The diffusion weighted images (DWI) were analyzed using the Tract-Based Spatial Statistics (TBSS) (Smith et al., 2006), part of FMRIB’s Software Library (FSL) (Smith et al., 2004). Image conversion from DICOM to NIfTI format was accomplished using dcm2nii. With FMRIB’s Diffusion Toolbox (FDT), the fractional anisotropy (FA) images were created by fitting a tensor model to the raw diffusion data and then, using the Brain Extraction Tool (BET) (Smith, 2002), brain-extracted. FA identifies the degree of anisotropy of a diffusion process and it is a measure often used in diffusion imaging where it is thought to reflect fiber density, axonal diameter, and myelination in white matter. A value of zero means that diffusion is isotropic, i.e. it is unrestricted (or equally restricted) in all directions, while a value of one means that diffusion occurs only along one axis and is fully restricted along all other directions. Subsequently the FA images were transformed into a common space by nonlinear registration IRTK(Rueckert et al., 1999). A mean FA skeleton, representing the centers of all tracts common to the group, was obtained from the thinned mean FA image. All FA data were projected onto this skeleton. The resulting data was fed into voxel-wise cross-subject statistics. Prior to analysis in SPM, the FA maps were converted from NIfTI format to Analyze.

The brains were segregated into 90 brain areas according to the Automated Anatomical Labeling Atlas 1 (AAL1) (Tzourio-Mazoyer et al., 2002). The anatomical names of the brain areas for each index *k* is shown in Tab. 1. In each brain network, one AAL brain area corresponds to a node of the network. The weights between the nodes were estimated through the measurement of the preferred diffusion directions, given by a set of *n_s_* = 5000 streamlines for each voxel. The streamlines are hypothesized to correlate with the white-matter tracts. The ratio of streamlines connecting area *l* and area *k* is given by the probability coefficient *p_lk_*. Then, the adjacency matrix *J_kl_* is constructed from this probability coefficient. The DTI processing pipeline has been adopted from Ref. (Cabral et al., 2013).

**Table 1.**
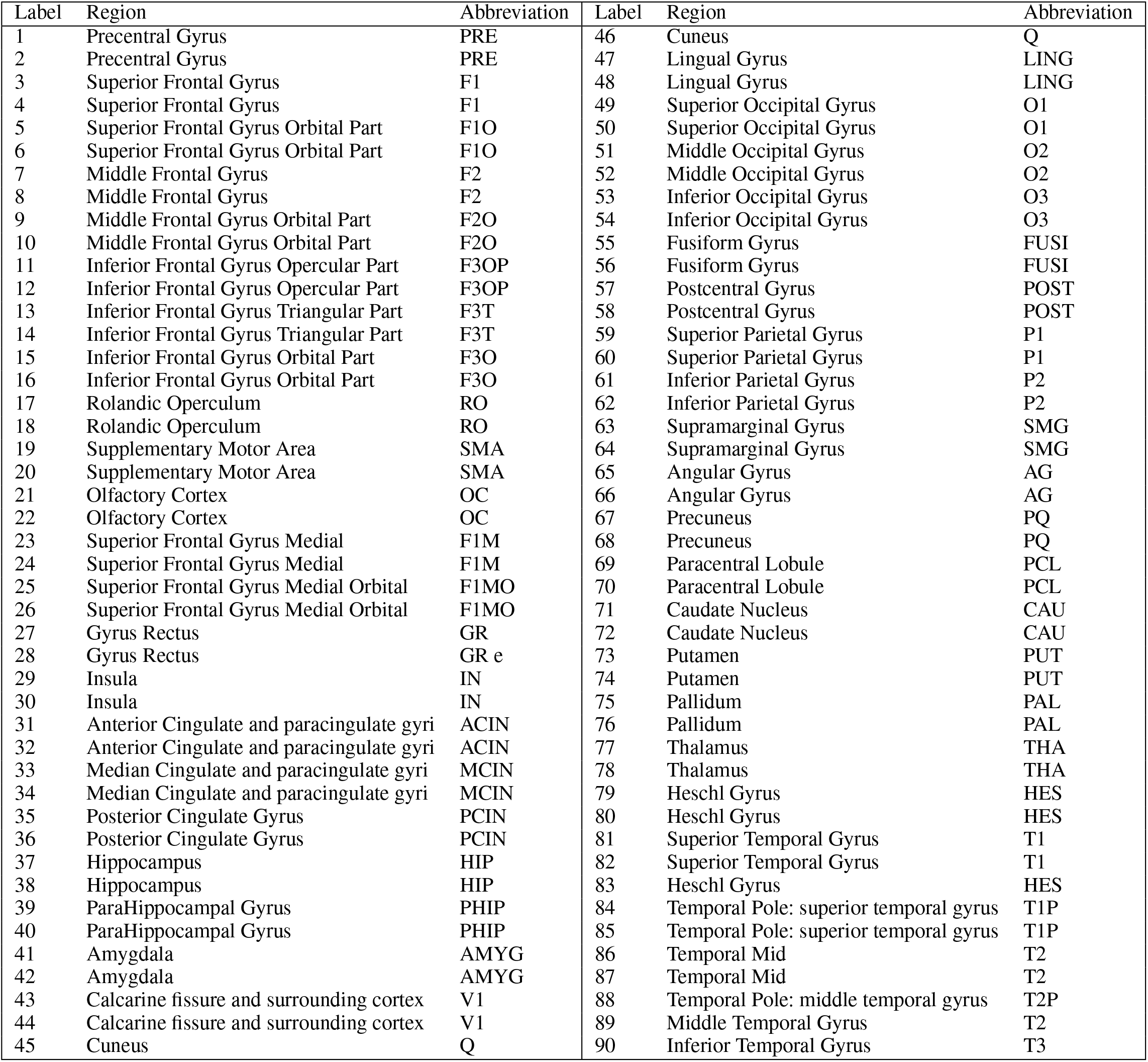
Cortical and subcortical regions, according to the Automated Anatomical Labeling atlas 1(AAL1) Tzourio-Mazoyer et al. (2002). Odd/even numbers correspond to the left/right hemisphere.

| Label | Region | Abbreviation | Label | Region | Abbreviation |
| --- | --- | --- | --- | --- | --- |
| 1 | Precentral Gyrus | PRE | 46 | Cuneus | Q |
| 2 | Precentral Gyrus | PRE | 47 | Lingual Gyrus | LING |
| 3 | Superior Frontal Gyrus | F1 | 48 | Lingual Gyrus | LING |
| 4 | Superior Frontal Gyrus | F1 | 49 | Superior Occipital Gyrus | O1 |
| 5 | Superior Frontal Gyrus Orbital Part | F1O | 50 | Superior Occipital Gyrus | O1 |
| 6 | Superior Frontal Gyrus Orbital Part | F1O | 51 | Middle Occipital Gyrus | O2 |
| 7 | Middle Frontal Gyrus | F2 | 52 | Middle Occipital Gyrus | O2 |
| 8 | Middle Frontal Gyrus | F2 | 53 | Inferior Occipital Gyrus | O3 |
| 9 | Middle Frontal Gyrus Orbital Part | F2O | 54 | Inferior Occipital Gyrus | O3 |
| 10 | Middle Frontal Gyrus Orbital Part | F2O | 55 | Fusiform Gyrus | FUSI |
| 11 | Inferior Frontal Gyrus Opercular Part | F3OP | 56 | Fusiform Gyrus | FUSI |
| 12 | Inferior Frontal Gyrus Opercular Part | F3OP | 57 | Postcentral Gyrus | POST |
| 13 | Inferior Frontal Gyrus Triangular Part | F3T | 58 | Postcentral Gyrus | POST |
| 14 | Inferior Frontal Gyrus Triangular Part | F3T | 59 | Superior Parietal Gyrus | P1 |
| 15 | Inferior Frontal Gyrus Orbital Part | F3O | 60 | Superior Parietal Gyrus | P1 |
| 16 | Inferior Frontal Gyrus Orbital Part | F3O | 61 | Inferior Parietal Gyrus | P2 |
| 17 | Rolandic Operculum | RO | 62 | Inferior Parietal Gyrus | P2 |
| 18 | Rolandic Operculum | RO | 63 | Supramarginal Gyrus | SMG |
| 19 | Supplementary Motor Area | SMA | 64 | Supramarginal Gyrus | SMG |
| 20 | Supplementary Motor Area | SMA | 65 | Angular Gyrus | AG |
| 21 | Olfactory Cortex | OC | 66 | Angular Gyrus | AG |
| 22 | Olfactory Cortex | OC | 67 | Precuneus | PQ |
| 23 | Superior Frontal Gyrus Medial | F1M | 68 | Precuneus | PQ |
| 24 | Superior Frontal Gyrus Medial | F1M | 69 | Paracentral Lobule | PCL |
| 25 | Superior Frontal Gyrus Medial Orbital | F1MO | 70 | Paracentral Lobule | PCL |
| 26 | Superior Frontal Gyrus Medial Orbital | F1MO | 71 | Caudate Nucleus | CAU |
| 27 | Gyrus Rectus | GR | 72 | Caudate Nucleus | CAU |
| 28 | Gyrus Rectus | GR e | 73 | Putamen | PUT |
| 29 | Insula | IN | 74 | Putamen | PUT |
| 30 | Insula | IN | 75 | Pallidum | PAL |
| 31 | Anterior Cingulate and paracingulate gyri | ACIN | 76 | Pallidum | PAL |
| 32 | Anterior Cingulate and paracingulate gyri | ACIN | 77 | Thalamus | THA |
| 33 | Median Cingulate and paracingulate gyri | MCIN | 78 | Thalamus | THA |
| 34 | Median Cingulate and paracingulate gyri | MCIN | 79 | Heschl Gyrus | HES |
| 35 | Posterior Cingulate Gyrus | PCIN | 80 | Heschl Gyrus | HES |
| 36 | Posterior Cingulate Gyrus | PCIN | 81 | Superior Temporal Gyrus | T1 |
| 37 | Hippocampus | HIP | 82 | Superior Temporal Gyrus | T1 |
| 38 | Hippocampus | HIP | 83 | Heschl Gyrus | HES |
| 39 | ParaHippocampal Gyrus | PHIP | 84 | Temporal Pole: superior temporal gyrus | T1P |
| 40 | ParaHippocampal Gyrus | PHIP | 85 | Temporal Pole: superior temporal gyrus | T1P |
| 41 | Amygdala | AMYG | 86 | Temporal Mid | T2 |
| 42 | Amygdala | AMYG | 87 | Temporal Mid | T2 |
| 43 | Calcarine fissure and surrounding cortex | V1 | 88 | Temporal Pole: middle temporal gyrus | T2P |
| 44 | Calcarine fissure and surrounding cortex | V1 | 89 | Middle Temporal Gyrus | T2 |
| 45 | Cuneus | Q | 90 | Inferior Temporal Gyrus | T3 |

Besides the healthy connectomes, we selected 15 connectomes (9 females, 6 males, mean age 33.4, range 22-56) of patients with different types of partial epilepsy that underwent a presurgical evaluation. The scans were performed at the Centre de Résonance Magnétique et Biologique et Médicale (Faculté de Médecine de la Timone) in Marseille. Diffusion MRI images were acquired on a Siemens Magnetom Verio 3T MR-scanner using a DTI-MR sequence with an angular gradient set of 64 directions, *TR* = 10700 ms, *TE* = 95 ms, 2 × 2 × 2*mm*^3^ voxel size, 70 slices, b-value 1000*s/mm*^2^.

The data processing pipeline (Schirner et al., 2015; Proix et al., 2016) made use of various tools such as FreeSurfer (Fischl, 2012), FSL (Jenkinson et al., 2012), MRtrix3 (Tournier, 2010) and Remesher (Fuhrmann et al., 2010), to reconstruct the individual cortical surface and large-scale connectivity. The surface was reconstructed using 20,000 vertices. Cortical and volumetric parcellations were performed using the Desikan-Killiany atlas with 70 cortical regions and 17 subcortical regions Desikan et al. (2006). The final atlas consists of 88 regions since one more empty region is added in the construction of the structural connectivity for symmetry. After correction of the diffusion data for eddy-currents and head motions using eddy-correct FSL functions, the Fiber orientation was estimated using Constrained Spherical Deconvolution (Tournier et al., 2007) and improved with Anatomically Constrained Tractography (Smith et al., 2012). For tractography, 2.5 × 10^6^ fibers were used and, for correction, Spherical-Deconvolution Informed Filtering of Tractograms (Smith et al., 2013) was applied. Summing track counts over each region of the parcellation yielded the adjacency matrix. Here, the AAL2 was employed for brain segregation leading to 88 brain areas for each patient, see Tab. 2.

**Table 2.**
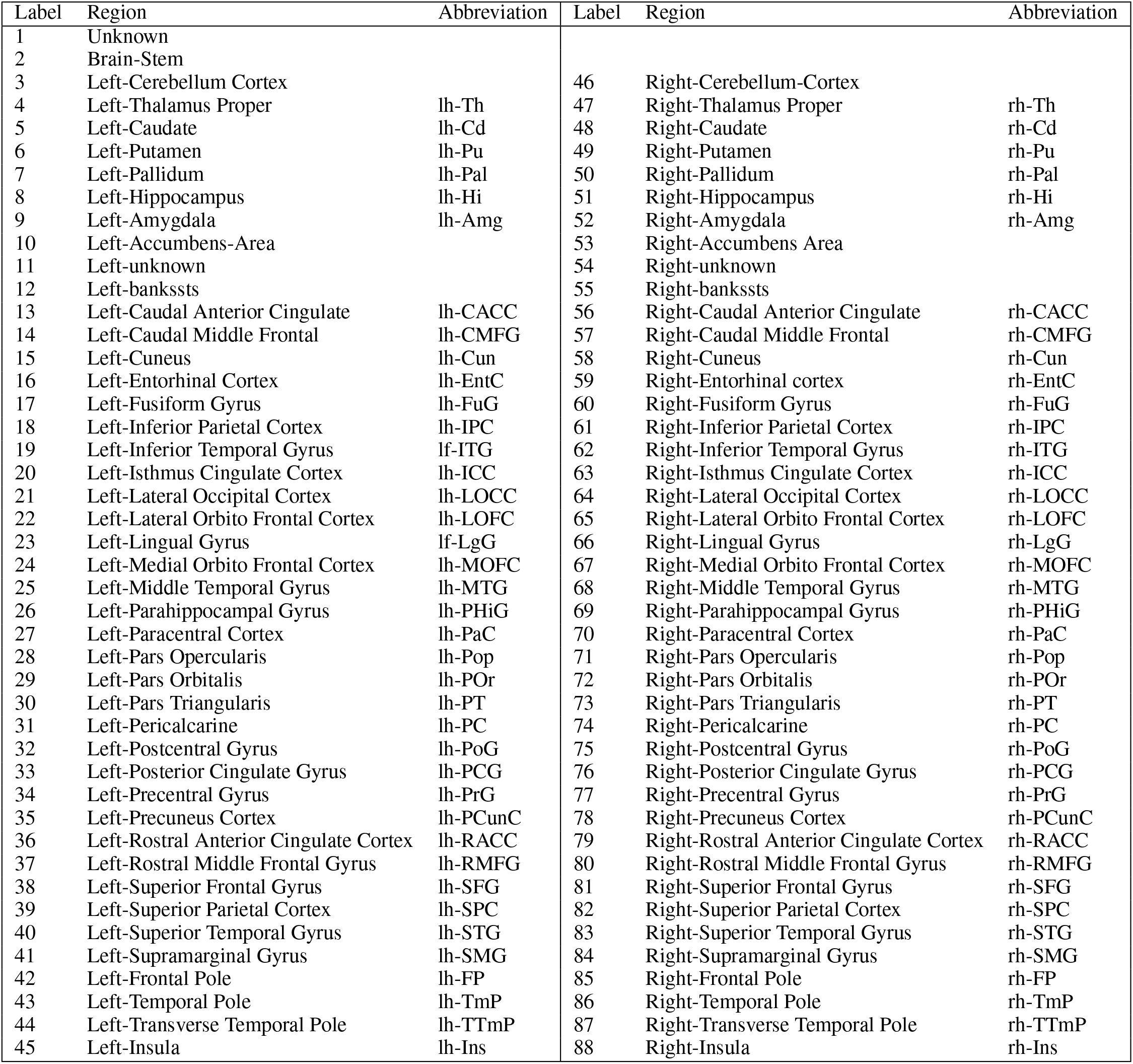
Cortical and subcortical regions, according to the Desikan-Killiany atlas Desikan et al. (2006).

### 2.5 EEG and SEEG data

All 15 drug-resistant patients, mentioned in the previous Section, affected by different types of partial epilepsy accounting for different Epileptogenic Zone (EZ) localizations, underwent a presurgical evaluation (see Supplementary Tables 3, 4). For each patient a first non-invasive evaluation procedure is foreseen, that comprises of the patient clinical record, neurological examinations, positron emission tomography (PET), and electroencephalography (EEG) along with video monitoring. Following this evaluation, potential EZs are identified by the clinicians. Further elaboration on the EZ is done in a second, invasive phase, which consists of positioning stereotactic EEG (SEEG) electrodes in or close to the suspected regions. These electrodes are equipped with 10 to 15 contacts that are 1.5 mm apart. Each contact has a length of 2 mm and measures 0.8 mm in diameter. Recordings were obtained using a 128 channel DeltamedTM system with a 256 Hz sampling rate and band-pass filtered between 0.16 Hz and 97 Hz by a hardware filter. All patients showed seizures in the SEEG, starting in one or several localized areas (EZ), before recruiting distant regions, identified as the Propagation Zone (PZ). Precise electrode positioning was performed by either a computerized tomography or MRI scan after implanting the electrodes.

Two methods were used for the identification of the propagation zone (see Supplementary Table 4). First, the clinicians evaluated the PZs subjectively on the basis of the EEG and SEEG recordings gathered throughout the two-step procedure (non-invasive and invasive). Second, the PZs were identified automatically based on the SEEG recordings: For each patient, all seizures were isolated in the SEEG time series. The bipolar SEEG was considered (between pairs of electrode contacts) and filtered between 1-50 Hz using a Butterworth band-pass filter. An area was defined as a PZ if its electrodes detected at least 30% of the maximum signal energy over all contacts, and if it was not in the EZ. In the following, we call the PZs identified by the subjective evaluation of clinicians PZ_Clin_ and the PZs identified through SEEG data PZ_SEEG_.

### 2.6 Network Measures

Topological properties of a network can be examined by using different graph measures that are provided by the general framework of the graph theory. These graph metrics can be classified in terms of measures that cover three main aspects of the topology: segregation, integration and centrality. The segregation accounts for the specialized processes that occur inside a restricted group of brain regions, usually densely connected, and it eventually reveals the presence of a dense neighborhood around a node, which results to be fundamental for the generation of clusters and cliques capable to share specialized information. Among the possible measures of segregation, we have considered the *clustering coefficient*, which gives the fraction of triangles around a node and it is equivalent to the fraction of node’s neighbors that are neighbors of each other as well. In particular the *average clustering coefficient C* of a network gives the fraction of closed triplets over the number of all open and closed triplets, where a triplet consists of three nodes with either two edges (open triplet) or three edges (closed triplet). The *weighted clustering coefficient* 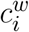 (Barrat et al., 2004) considers the weights of its neighbors:

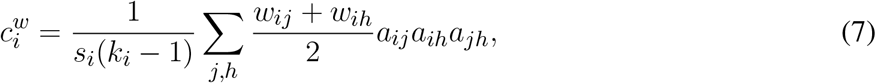

where *s_i_* is the node strength (to be defined below), *k_i_* the node degree, *w_ij_* the weight of the link, and *a_ij_* is 1 if the link *i* → *j* exists and 0 if node *i* and *j* are not connected. The *average weighted clustering coefficient C_W_* is the mean of all weighted clustering coefficients: 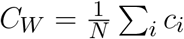.

The measures of integration refer to the capacity of the network to rapidly combine specialized information from not nearby, distributed regions. Integration measures are based on the concept of communication paths and path lengths, which estimate the unique sequence of nodes and links that are able to carry the transmission flow of information between pairs of brain regions. The *shortest path d_ij_* between two nodes is the path with the least number of links. The *average shortest path length* of node *i* of a graph *G* is the mean of all shortest paths from node *i* to all other nodes of the network: 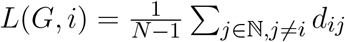. The *average shortest path length* of all nodes is the mean of all shortest paths (Boccaletti et al., 2006): 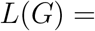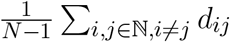. In a weighted network, distance and weight have a reciprocal relation. If a weight between two adjacent nodes is doubled, their shortest path is cut by half: 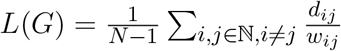.

Centrality refers to the importance of network nodes and edges for the network functioning. The most intuitive index of centrality is the node degree, which gives the number of links connected to the node; for this measure, connection weights are ignored in calculations. In this manuscript, we employ the network measure *node strength s_i_*, which corresponds to the weighted node degree of node *i* and equals the sum of all its weights: *s_i_* = ∑_*j*∈ℕ_ *w_ij_*. Accordingly, the *average node strength S* corresponds to the mean of all node strengths 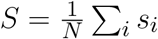. All finite networks have a finite number of shortest paths *d*(*i, j*) between any pair of nodes *i*, *j*. The *betweenness centrality c_B_*(*s*) of node *s* is equal to all pairs of shortest paths that pass through *s* divided by the number of all shortest paths in the network: 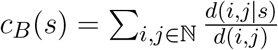. For the *weighted betweenness centrality*, the *weighted shorted paths* are used.

## 3 RESULTS

The detection of epileptic seizures via electrophysiological recordings allowed for the establishment of a detailed taxonomy of seizures. The majority of seizures recorded in humans and experimental animal models can be described by a generic phenomenological mathematical model, the Epileptor Jirsa et al. (2014). In this model, seizure events are driven by a slow permittivity variable and occur via saddle node and homoclinic bifurcations at seizure onset and offset, respectively. The saddle-node bifurcation at the onset of ictal discharges was chosen based on experimentally observed features, such as fixed frequency and fixed amplitude of abruptly starting oscillations, and a shift of baseline field potential. The homoclinic bifurcation at the offset of ictal discharges, on the other hand, reproduces the logarithmic scaling of interspike intervals when approaching seizure offset. As part of the dynamic repertoire of the Epileptor, the epileptic attractor is described in terms of a self-sustained limit cycle that comes from the destabilisation of the physiological activity while multiple types of transitions allow for the accessibility of seizure activity, status epilepticus and depolarization block, that coexist, as verified experimentally in El Houssaini et al. (2020).

The Epileptor model has been reduced to a minimal canonical mathematical representation of high codimension (up to 4) that, appropriately tuned, can display several types of fast-slow behaviors Saggio et al. (2017). The model contains two subsystems acting at different time scales, in which the fast subsystem is unfolded in a plane showing several bifurcation paths of a high codimension singularity. The slow subsystem steers the fast one back and forth along these paths leading to fast-slow (aka bursting) behavior, mimicking epileptiform activity. The model is able to produce almost all the classes of bursting predicted for systems with a planar fast subsystem, including the Epileptor class, which is also the target class in this paper and has been demonstrated to be the dominant class, so-called dynamotype, in empirical epilepsy data Saggio et al. (2020). Other dynamotypes have been also found empirically.

When performing the analysis of the single-population firing rate equations (4), it turns out that, in the absence of forcing, the only attractors are fixed points. As it will become clear in the following Section, a stable node and a stable focus are observable, separated by a bistability region between a high- and a low-activity state, whose boundaries are the locus of a saddle-node bifurcation (for more details see (Montbrió et al., 2015)). In this context are not observable self-sustained oscillations, but only damped oscillations at the macroscopic level that reflect the oscillatory decay to the stable fixed point. This oscillatory decay will here be considered as representative of a seizure-like event, not being able to observe a stable limit cycle to describe the emergence of a fully developed seizure as in the Epileptor. However, seizure-like events can be used as paradigm to investigate propagation of seizure-like activity in the network. Furthermore, a recently developed model of interictal and ictal discharges, called Epileptor-2 Chizhov et al. (2018), makes links to underlying physiology and suggest how to eventually obtain all observed dynamotypes for the exact neural mass model (4) and enable transitions towards fully developed seizure activity.

Epileptor-2 is a simple population-type model that includes four principal variables, i.e. the extracellular potassium concentration, the intracellular sodium concentration, the membrane potential and the synaptic resource diminishing due to short-term synaptic depression. A QIF neuron model, whose dynamics is ruled by an equation similar to Eq. (1), is used as an observer of the population activity. While the potassium accumulation governs the transition from the silent state to the state of ictal discharge, the sodium accumulated during the discharge, activates the sodium-potassium pump, which terminates the ictal discharge by restoring the potassium gradient, thus polarizing the neuronal membranes. This means that, in high potassium conditions, Epileptor-2 produces bursts of bursts, described as ictal-like discharges.

Therefore, the association of a slow subsystem describing ion concentration variations together with a fast subsystem, identified by Eqs. (4), should give rise to self-emergent periodic and bursting dynamics at the macroscopic level, thus allowing us to identify different combinations of onset/offset bifurcations. Whenever not sufficient, it will be possible to investigate the dynamics emergent in the exact neural mass model, provided with short-term synaptic plasticity, when subject to a global feedback acting on a slow timescale, describing ion concentration variations. The exact neural mass model, when equipped with short-term synaptic plasticity, shows a more complex dynamics that eventually results in a bifurcation diagram that provides stable limit cycles Taher et al. (2020). However the introduction of short-term plasticity, itself, adds complexity to the dynamics, allowing for the emergence of bursting activity Tsodyks et al. (1998).

### 3.1 Healthy Subjects

#### 3.1.1 Phase and Bifurcation Diagrams

The analysis of the single-population firing rate equations (4), performed in (Montbrió et al., 2015), has revealed that there are three distinct regions, when considering the phase diagram of the system as a function of the external drive 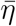 and synaptic weight *J*, in absence of time dependent forcing (*I*(*t*) = 0): (1) a single stable node equilibrium corresponding to a low-activity state, (2) a single stable focus (spiral) generally corresponding to a high-activity state, and (3) a region of bistability between low and high firing rate. In particular, in the region where the stable focus is observable, the system undergoes damped oscillatory motion towards this fixed point. The presence of damped oscillations at the macroscopic level reflects the transitory synchronous firing of a fraction of the neurons in the ensemble. While this behavior is common in network models of spiking neurons, it is not captured by traditional firing-rate models (Schaffer et al., 2013; Devalle et al., 2017; Taher et al., 2020).

When considering the multipopulation neural mass model (5) with homogeneously set 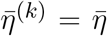, the corresponding phase diagram (shown in Fig. 1 B) is qualitatively the same as the one shown in Fig 1 in (Montbrió et al., 2015), since the same attractors are observable. In the original model these attractors are single-population states, while they reflect multipopulation states in the present case. Single-population low-activity (LA) and high-activity (HA) states translate into network LA and HA states. In the former all populations have low, in the latter high firing rates. We observe that the single-population bistability accurately reflects the hysteretic transition in the network when changing 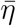. In the following we will address how this relation between single-node and multipopulation phase diagram occurs.

**Figure 1.**
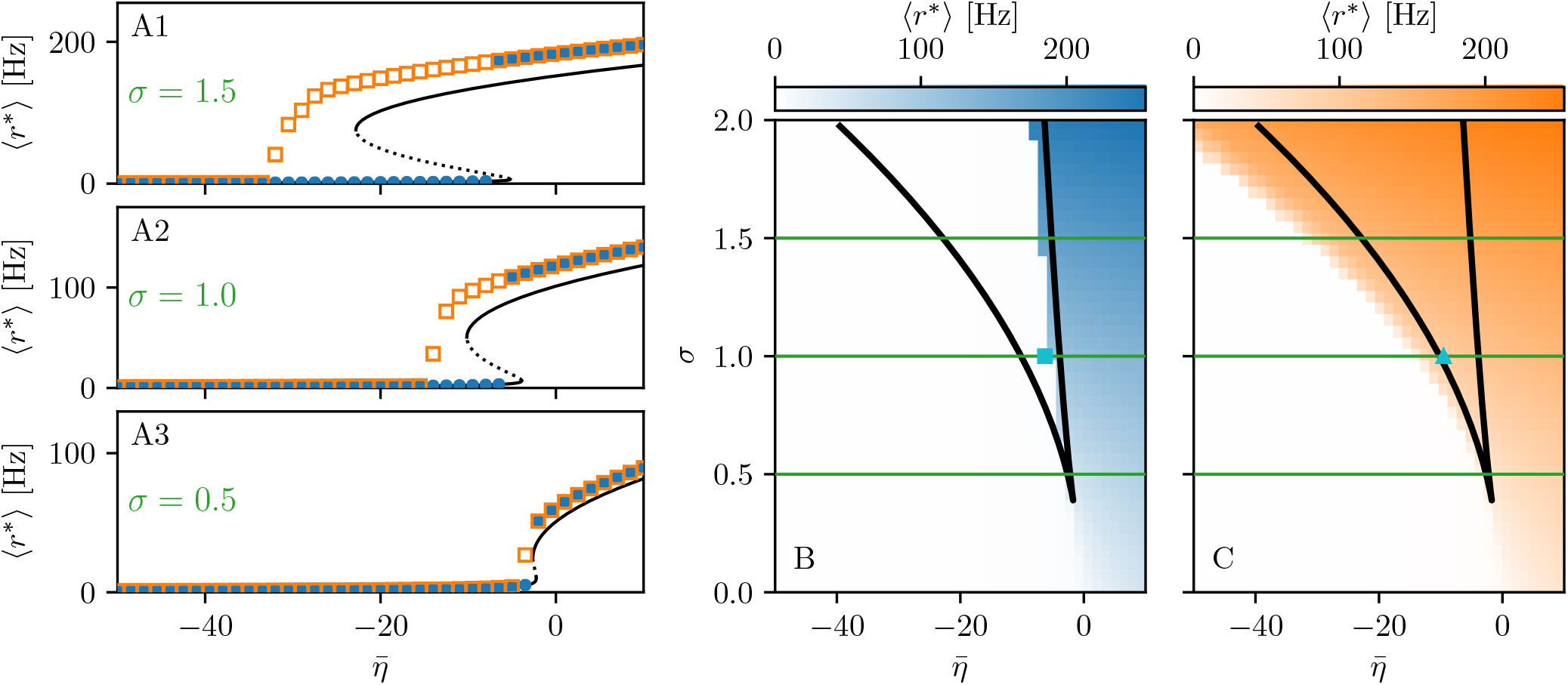
A1-A3 Equilibrium firing rates 〈*r*\*〉 vs. 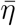 for the up-sweep (blue dots) and down-sweep (orange squares). For each 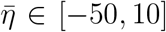 in steps of 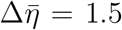 the system is initialized using the final state of the previous run and evolves for 2 s after which the average network firing rate in the equilibrium state is determined. Different panels correspond to different *σ* values: *σ* = 1.5 (A1), *σ* = 1 (A2), *σ* = 0.5 (A3). The solid (dashed) black line corresponds to the stable (unstable) equilibria in the single-node case. Maps of regimes as a function of *σ* and 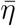 showing the network average 〈*r*\*〉 color coded for up- (B) and down-sweep (C), obtained by following the same procedure as in A1-A3 for *σ* ∈ [0, 2] in steps of Δ*σ* = 0.05. The black line indicates the single-node map of regimes like in (Montbrio et al., 2015). In panels B-C the cyan square and triangle mark 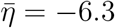, −9.54 respectively. Parameter values: *N*_pop_ = 90, *τ*_m_ = 20 ms, Δ = 1, *J_kk_* = 20, 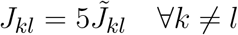.

The network bifurcation diagrams shown in panels A1-A3 for increasing *σ* values are obtained by performing an adiabatic analysis along two different protocols: up-sweep and down-sweep. Following the up-sweep protocol, the system’s state variables *r_k_, v_k_* are initialized at 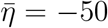 with the values *r_k_* = 0, *v_k_* = 0; then the excitability is increased in steps 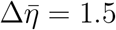 until the maximal value 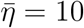 is reached. At each step, the initial conditions for mean firing rates and mean membrane potentials correspond to the final state obtained for the previous 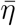 value. Note, that the average firing rate increases for increasing 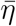 values, both for the single node and for the network. Once the maximum 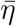 value is reached, the reverse procedure is performed, thus following the down-sweep protocol. This time the initial conditions correspond to the high-activity state at 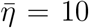, while the excitability is adiabatically decreased in steps 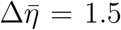, until a low-activity state at 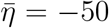 is approached. For both protocols, the investigation of the nature of the dynamics emerging at each 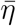-step is done by using the same procedure: the system is simulated for a transient time *T* = 2 s, until it has reached an equilibrium state. At this time the firing rate averaged over all populations 〈*r*\*〉 is calculated and the next 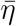 iteration is started, using this final state as initial conditions.

The transition from LA to HA network dynamics is hysteretic: the system doesn’t follow the same path during the up-sweep and the down-sweep protocol. When the system is initialized in the low activity regime, it remains there until a critical excitability value 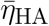 is reached. For further increase of the excitability, the average firing rate exhibits a rapid jump to higher values. However, when the system is initialized in the high-activity regime, this regime survives for a large 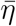 interval until it collapses toward a low-activity state at 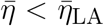, where 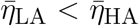. There is a considerable difference between the critical excitability values required to lead the system to a high-activity or a low-activity regime and the difference increases for increasing coupling strength *σ*. While the up-sweep protocol (blue dots) is well approximated by the bifurcation diagram of the single population, represented in panels A1-A3 by the black (dashed and continuous) curve, this is no more true for the down-sweep protocol, where the coupling plays a role in determining the transition at the multipopulation level (orange squares). This results in different phase diagrams for the two protocols: the maps of regimes is dominated by the low-activity (high-activity) state when following the up-sweep (down-sweep) protocol. Merging together these results we observe that the region of bistability between LA and HA network dynamics, is still identifiable by the original boundaries found for the single population in (Montbrió et al., 2015) (see black curve in panels B, C), even though, for the multipopulation system, the region is wider.

We can make further use of the single-population bifurcation diagram to understand the hysteretic transition of the multipopulation model in more detail. First of all, the weight matrix {*J_kl_*} has its largest components on the diagonal (*J_kk_* = 20), reflecting recurrent synapses. This means that synaptic inter-coupling plays a minor role, as long as the firing rates of the adjacent populations are small. During the up-sweep protocol, this condition is fulfilled, as all populations are initialized in a low activity regime. Under these conditions, the dynamics of all nodes is rendered identical and equal, approximately, to the single population dynamics. Consequently the single-population LA branch describes the multipopulation LA behaviour (in terms of 〈*r*\*〉) accurately as a function of 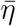. Secondly, as soon as the single-population LA state vanishes for large enough 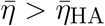, the individual nodes of the multipopulation system all transit to the HA state.

In this HA regime, deviations of the network bifurcation diagram with respect to the single-population curve are observed. The populations in the system have large firing rates, such that the inter-coupling becomes a relevant contribution to the total current on each node. This explains why the LA branch of the network is located at higher firing rates with respect to the black single-population curve: The populations in the network behave, approximately, as decoupled, irrespectively of being subject, in the HA regime, to an additional current due to the inter-coupling. This effectively shifts the single-population bifurcation diagram towards smaller 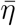. Moreover this shift occurs for each population individually, depending on the matrix {*J_kl_*}. During the down-sweep protocol, due to the population dependent shift, the HA population states vanish at different values of 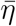. Accordingly, whenever this occurs, the network average 〈*r*\*〉 decreases by a small amount, such that the network LA state is reached via various intermediate states. We can infer, using the same type of argument, that single-population LA states disappear for increasing 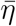 in a region around 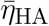. They are not observed here, due to the nature of the up-sweep protocol and the initialization procedure of *r_k_, v_k_*.

From the reversed viewpoint we can hypothesize, that stable single-population HA states may take form near 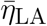 for increasing 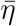, as well as stable LA states for decreasing 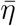 near 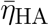. This implies that the network possesses complex multistability between many network states in the region 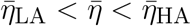. For these states the existence of LA and HA states of individual populations are interdependent: whether or not any given population can be in the LA or HA state is conditioned by the LA-HA configuration of all other populations. This not only demonstrates how multistability emerges in the multipopulation system, but it also has influence on the response of the network towards transient input in such a setting. Most importantly, if such an input recruits one population into high activity, other populations might follow, leading to a cascade of recruitments.

#### 3.1.2 Seizure-like Recruitment in Dependence of Perturbation Site and 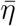

To analyze the response of the multipopulation system to transient current, we stimulate one population with a step function *I_S_*(*t*) of amplitude *I_S_* = 10 and duration *t_I_* = 0.4 s. By setting 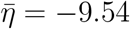, the system is placed in the multistable regime (see cyan triangle in Fig. 1C), but, due to the low 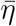 value, it only allows for LA-HA configurations with most of the populations in LA. We start by initializing all nodes in the low-activity state and stimulating a single node (see Fig. 2 A). During the stimulation (panel A1), the stable states of the network change, due to the strong additional current. More specifically, the initial equilibrium vanishes and a new focus equilibrium of the system appears as the only stable network state. This focus is characterized by an LA-HA configuration for which only the stimulated node finds itself in HA while the rest remains in the LA regime; the focus is approached via damped oscillations in the time interval 0 *< t <* 0.4 s (panel A2-A3). Due to the multistability in absence of stimulation, an identical LA-HA configuration exists. Thus, when the current is removed, the system is able to maintain the LA-HA configuration. However, the position of the focus equilibrium is shifted in absence of the transient input and is reached, again, via damped oscillations for *t >* 0.4 s.

**Figure 2.**
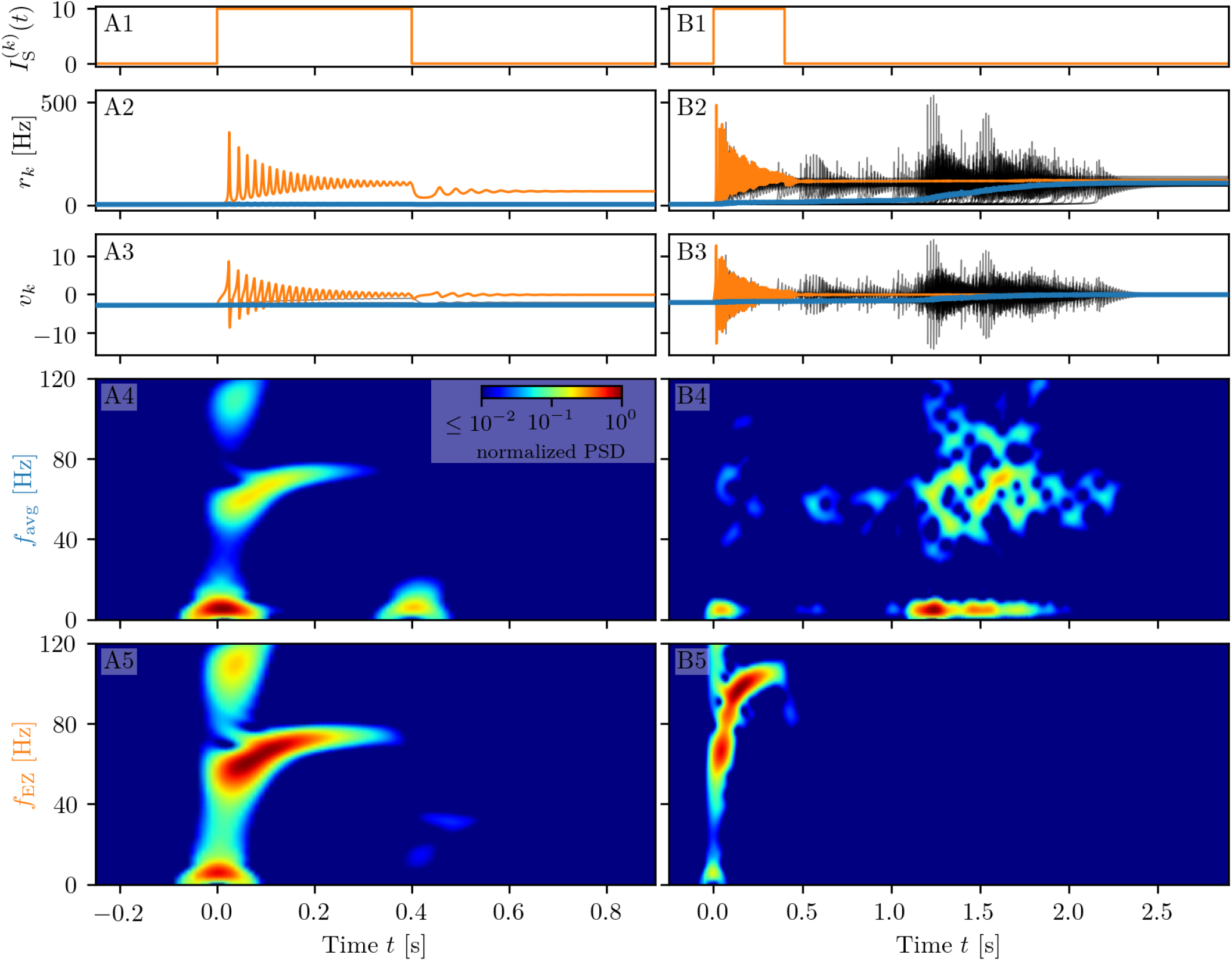
Spectrograms of mean membrane potentials for subject sc0. (A1-B1) Stimulation current 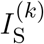, (A2-B2) population firing rates *r_k_* and (A3-B3) mean membrane potentials *v_k_* for the EZ (orange) and other populations (black). The blue curves show the network average firing rate and membrane potential. Non-stimulated node dynamics is plotted as transparent gray curves: some of the nodes adapt their voltage to the stimulation of the EZ and change during stimulation. However they do not reach the high-activity state regime. (A4-B4) Spectrogram of the network average membrane potential and (A5-B5) of the *v_k_* of the EZ. Column A shows an asymptomatic seizure-like event for 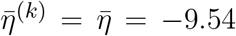, column B a generalized seizure-like event for 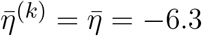. In both cases the EZ node 46 is stimulated. Parameter values: *N*_pop_ = 90, *τ*_m_ = 20 ms, Δ = 1, *J_kk_* = 20, *σ* = 1, 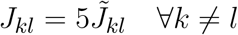.

When the perturbation of a single node has no consequences on the dynamics of the other populations, as shown in Fig. 2 A2), A3), we are in the presence of an *asymptomatic seizure-like event*, where the activity is limited to the epileptogenic zone (here represented by the stimulated node) and no propagation takes place. For higher excitability values (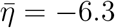, marked by a cyan rectangle in Fig. 1B), the perturbation of a single node gives rise to a different response dynamics. In this case other brain areas are “recruited” and not only the perturbed node, but also other populations reach the high-activity regime by showing damped oscillations (see Fig. 2 panels B2, B3). In terms of pathological activity, the seizure-like event originates in the EZ (as a results of the stimulation) and propagates to the PZ, identified by the other regions which rapidly propagate the oscillatory activity. The recruitment of the regions in the propagation zone can happen either by independent activation of the single areas, or by activating multiple areas at the same time, until the propagation involves almost all populations (*generalized seizure-like event*).

The transition of a single population to the HA regime, upon stimulus onset, is characterized by a transient activity in the *δ* band (*<* 12 Hz) and a sustained activity in the *γ* band (40-80 Hz), present throughout the stimulation, as shown in Fig. 2, panels A4-A5. Here the spectrograms show time varying power spectral densities (PSD) of the mean membrane potentials averaged over the network (A4) and for the single stimulated population (A5). When more populations are recruited at higher excitability values, in addition to the former activity, it is possible to observe *γ* activity at higher frequencies (see panels B4-B5). High-frequency oscillations, between 80 and 500 Hz, can be recorded with EEG and reflect the seizure-generating capability of the underlying tissue, thus being used as markers of the epileptogenic zone (Jacobs et al., 2012). Moreover, even for the generalized seizure-like case, the *δ* band activity is evoked whenever a brain area gets recruited, leading to a sustained signal in the time interval 1.1 s *< t <* 1.8s, where a majority of the populations approach the HA state. Similar results have been obtained for all the other investigated subjects (results not shown).

In the following we report a wide analysis of the impact of the perturbation site on the recruitment effect, for different excitability values. As before, we use a step current *I*_S_(*t*), with fixed amplitude *I_S_* = 10 and duration *t_I_* = 0.4 s, to excite a single population. In each run the stimulating current targets a different brain area and the number of recruitments, i.e. the number of populations, that pass from the LA state to the HA state, is counted. The 90 brain areas are targeted, one at a time, in 90 individual simulations. We repeat the procedure varying 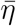 in a range [−15, −4], with steps of 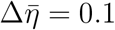. The results for five exemplary subjects are shown in Fig. 3 A1)-E1).

**Figure 3.**
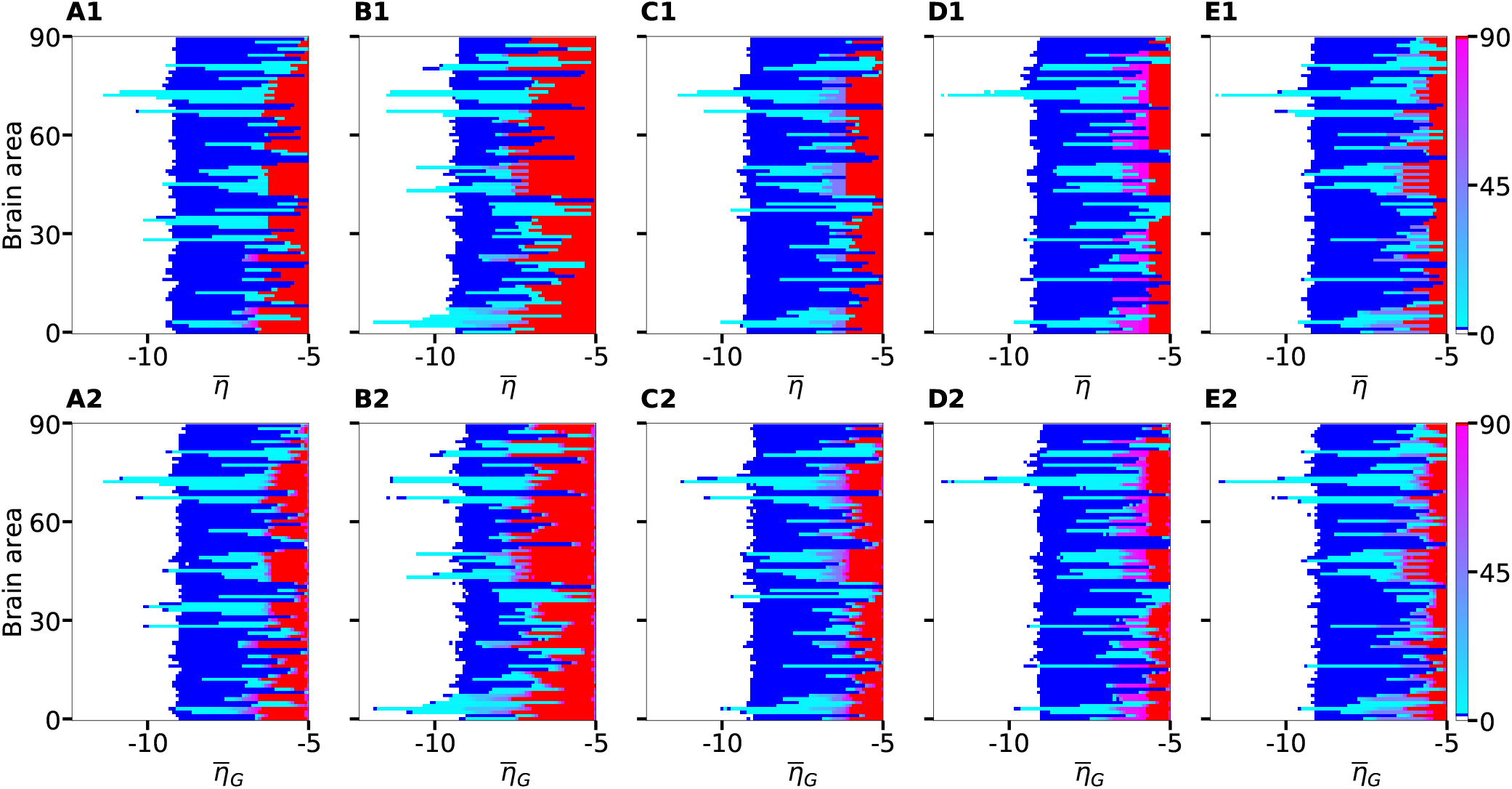
Number of recruited brain areas as a function of the excitability parameter 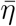 for 5 exemplary healthy subject connectomes A-E. Color coding is the following: blue corresponds to the asymptomatic threshold (one area in HA regime); red represents 90 areas in HA regime (generalized threshold); cyan to purple indicate intermediate recruitment values, white marks no recruitment. When performing a vertical cut, all nodes are characterized by the same 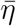 for panels (A1-E1). At the contrary, in panels (A2-E2), 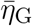 represents the mean value of a Gaussian distribution with standard deviation 0.1. Therefore, when perturbing one brain area at a time, excitabilities are distributed and not uniform in the latter case; the results are averaged over 10 repetitions with different Gaussian excitability distributions. A), B), C), D), and E) correspond to subjects 0, 4, 11, 15, and 18. Parameters: *N* = 90, Δ = 1, *σ* = 1, *I_S_* = 10, *t_I_* = 0.4 s.

If the perturbed area jumps back to the LA state when the stimulation is removed and no further recruitment takes place, then the total number of recruited areas is zero, here color coded in white. If the perturbed area remains in the HA state without recruiting other areas, we are in presence of an asymptomatic seizure-like event (blue regions). For every further recruited brain area, the color code changes from cyan to purple. If all brain areas are recruited, we observe a generalized seizure-like event (coded as red). For 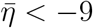, most of the targeted brain areas goes back to the LA state, when the perturbation ends, while for 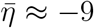, we generally observe asymptomatic seizure-like events for all the subjects and for most of the perturbation sites. For increasing 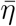 values, the probability for larger recruitment cascades increases, until the system exhibits generalized seizure-like events for 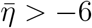. However, some notable differences between brain areas and among the different subjects are observable. Brain area 72, for example, corresponding to the rh-CAU, exhibits asymptomatic seizure-like events at 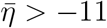 for most of the subjects, thus suggesting that the rh-CAU favours pathological behavior with respect to other brain areas. On the other hand, some brain areas are less likely to cause generalized seizure-like events, when stimulated, than others: Brain area 40, for example, the rh-PHIP^1^, causes no generalized seizure-like events for any 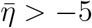. Note that, for very large 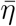 values, the system doesn’t exhibit multistability anymore, but instead has only one stable state, namely the network HA state, corresponding to high firing rate of all populations. Approximately, this happens for 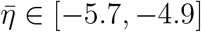, with small variations among the subjects.

The scenario remains unchanged when we take into account heterogeneous excitabilities 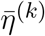, as shown in Fig. 3 A2)-E2). In this case 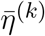 is drawn from a Gaussian distribution with mean 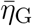, thus mimicking the variability present in a real brain. The populations are stimulated, as before, one at a time in individual simulation runs. However, this time the procedure is repeated for varying 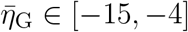, while keeping the standard deviation of the Gaussian distribution fixed at 0.1. Larger standard deviations hinder the rich multistability of the network, by eliminating the bistability between LA and HA for individual populations, due to excessively small or large 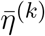, thus impeding the analysis of the impact of the stimulation. The shown results are obtained averaging over 10 Gaussian distribution realizations of the 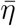 parameter; slightly more variability becomes apparent especially when considering the threshold in 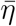 to observe generalized seizures.

An overview over all the investigated subjects is possible when looking at Fig. 4 A), where is reported the average, over all subjects, of the data shown in Fig. 3 A1)-E1) for five exemplary subjects only. The averaging operation smears out the transition contours and, while the region of generalized seizure-like events shrinks, it becomes wider the region of accessibility of partial seizure-like events, where a small percentage of nodes (~ 20%) are recruited. In panel B we report 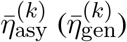, i.e. the smallest 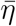 value for which an asymptomatic (generalized) seizure-like event occurs when stimulating population *k*. Grey dots indicate the individual thresholds 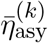 and 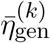 for each of the 20 subjects and 90 brain areas; the averages over all subjects are denoted by blue and red circles, respectively, for each *k* ∈ [1, 90]. Averaging these thresholds over all subjects and brain areas yields an asymptotic threshold of 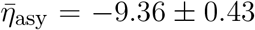 and a generalized threshold of 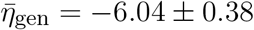. Brain areas 72, 73, 67, and 3 have lower thresholds for asymptomatic seizure-like events, areas 40, 86, and 82 have larger thresholds for generalized seizure-like events and do not fall within a standard deviation. The variability in the response among the different areas is more evident for 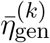 values compared to the 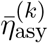 ones: the threshold values to obtain an asymptomatic seizure-like event are very similar among the areas and among the subjects, while the threshold values to obtain a generalized seizure-like event strongly depend on the stimulated area and on the subject.

**Figure 4.**
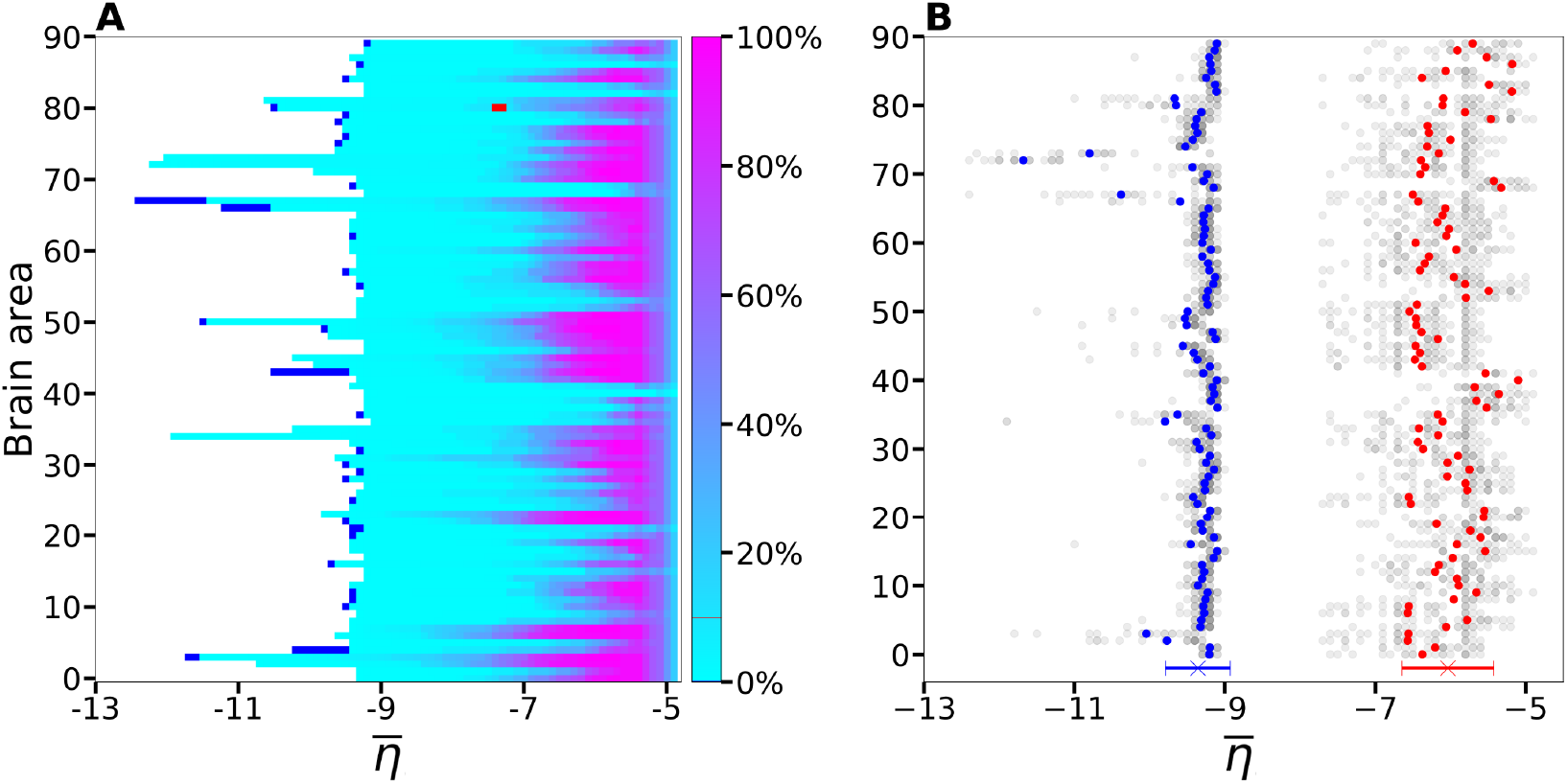
A) Number of recruited brain areas as a function of the excitability parameter 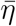, as shown in Fig. 3 A1)-E1), averaged across all subjects. B) 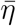 threshold values for asymptomatic and generalized seizure-like events. Grey dots show the thresholds for each brain area and each subject. Blue and red dots show the average over 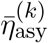 and 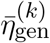 across all subjects. The blue and red cross at the bottom show the average value and its standard deviation for both thresholds across all subjects and across all areas. Parameters as in Fig. 3.

#### 3.1.3 The Role Played by Brain Area Network Measures on Enhancing Recruitment

As shown in Fig. 4 B), 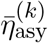 does not vary significantly among the subjects and among the brain areas; it mainly occurs in the range 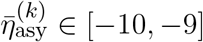, with just few nodes (*k* ∈ [72, 73, 67, 3]) showing smaller values. Since each brain area is characterized by its own network measure, the first hypothesis that we aim to test, is the role played, on the identification of the threshold, by the different network measures. In particular, we investigate the dependency of 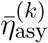 on the node strength, clustering coefficient, shortest path length, and betweenness centrality of the corresponding brain area, as shown in Fig. 5. A very strong correlation between asymptomatic threshold and node strength becomes apparent: Brain areas that are strongly connected, need a smaller excitability to pass from the LA to the HA regime (panel A). The same holds true for the clustering coefficient, even though the relationship is less sharp (panel B). Moreover it is possible to observe a direct correlation between 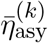 and shortest path length (i.e. shortest is the path smallest is the threshold value), while betweenness is smaller for higher threshold values (panels C and D respectively).

**Figure 5.**
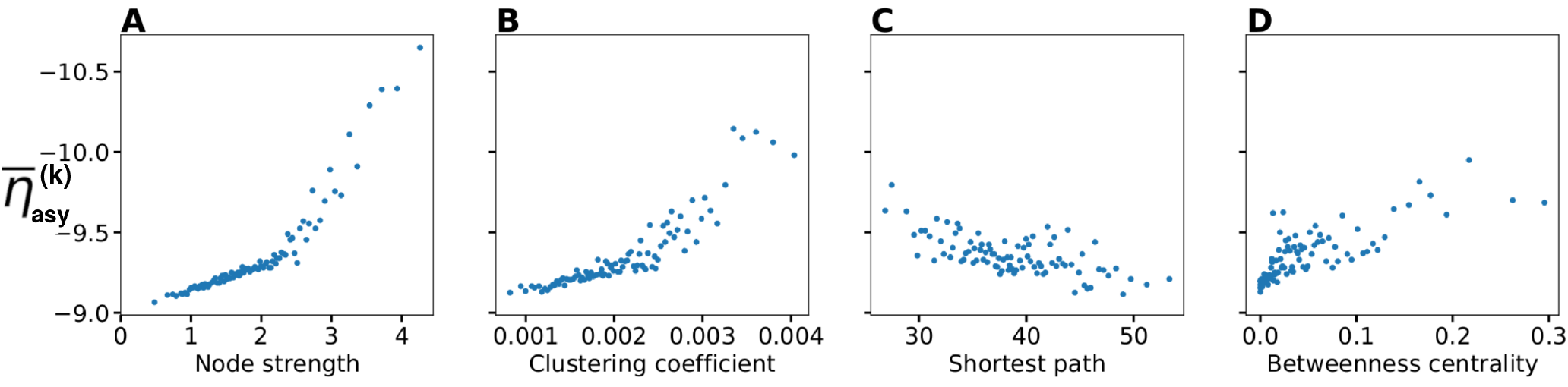
Threshold 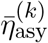 for asymptomatic seizure-like events as a function of node measures: A) Node strength, B) clustering coefficient, C) average shortest path length, D) betweenness centrality. For each panel, the thresholds 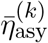 are calculated for all *k* ∈ [1, 90] brain areas and averaged over all 20 subjects. Parameters as in Fig. 3.

When considering the threshold for generalized seizure-like events, we face a higher variability among different nodes (as shown in Fig. 4B, 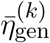 varies mainly between −6.5 and −5.5). The dependency of 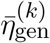 on the node strength reveals a strong correlation: Areas with very small node strengths are characterized by large thresholds and are less likely to cause generalized seizure-like events. On the other hand, for large node strengths, 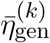 saturates at a value ≈ −6.5 (see Fig. 6 A)). The clustering coefficient, shown in Fig. 6 B), shows a similar relationship as the node strength, even though more scattered. This is not surprising since node strength and clustering coefficient are strongly correlated with each other (the Pearson Correlation coefficient in this case is *r* = 0.75, as shown in Fig. 1 of the Supplementary Material), thus explaining the similarity between the analyses reported in panels A) and B). Moreover, regarding the integration measure, it turns out that the average shortest path length correlates positively with 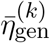 (see Fig. 6 C)). Brain areas that are characterized, on average, by a short path to all the other areas are more likely to cause generalized seizure-like events. Finally, the betweenness centrality correlates negatively with 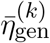 (panel D). This means that brain areas that are crossed by many shortest path lengths (large betweenness centrality) are more likely to cause generalized seizure-like events. For increasing node strength, clustering coefficient and betweenness centrality, we observe a saturation toward 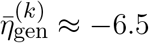, that corresponds to the critical excitability value, during the up-sweep simulation, at which the system jumps to the HA network state (Fig. 1 A2).

**Figure 6.**
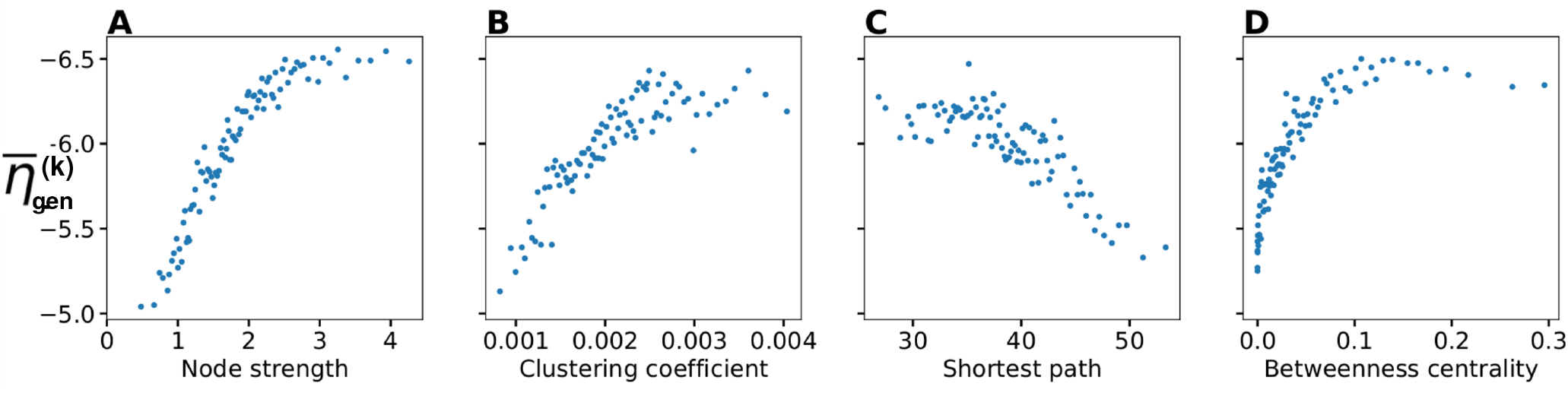
Threshold for generalized seizure-like events as a function of node measures: A) Node strength, B) clustering coefficient, C) average shortest path length, D) betweenness centrality. For each panel, the thresholds 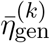 are calculated for all *k* ∈ [1, 90] brain areas and averaged over all 20 subjects. Parameters as in Fig. 3.

To explore the causal mechanisms of brain function and understand the sequential mechanism of node recruitment in more detail, we investigate the timing at which different brain areas are recruited. For this, the excitability parameter 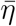, common to all populations, is set to the threshold value 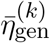 of the perturbed brain area *k*, ensuring complete recruitment of all populations, when perturbing populations *k* ∈ [1, 90]. The results shown in Fig. 7 are obtained by averaging over *k* and over the different subjects: in 90 individual simulations for each subject, a single brain area *k* = 1, …, 90 is stimulated with an external step current *I*_S_(*t*), characterized by an amplitude *I*_S_ = 10 and a duration *t*_I_ = 0.4 s. For each *k* the recruitment time of all the other areas is registered. The stimulated brain area stands in for the EZ. The brain areas and corresponding node measures are sorted by the recruitment time in ascending order. The weight and shortest path values taken into account are divided in two types: those related to the nodes outgoing the EZ (i.e. the subgraph connected to the EZ), and those related to the connections between the recruited area and all the other nodes except the EZ. Therefore, for the first case, if a certain recruited area is not directly connected to the EZ, its corresponding weight is equal to 0. The values for recruitment time (panel A), weight of a connection between a single area and the EZ (panel B) and shortest path (panel C) are finally obtained averaging over all the stimulated nodes and all the subjects (i.e. the average is performed over 1800 simulations across all 90 brain area perturbations times all 20 subject). The same averaging procedure has been employed to obtain the data shown in panels D-G. However, in this case, the node measures are evaluated over all the connections of the recruited node minus the connection to the EZ. While ignoring the link to the excited area (EZ), the overall network measure for connection weights (panel D), clustering coefficient (panel E), shortest path (panel F) and betweenness centrality (panel G) are reported.

**Figure 7.**
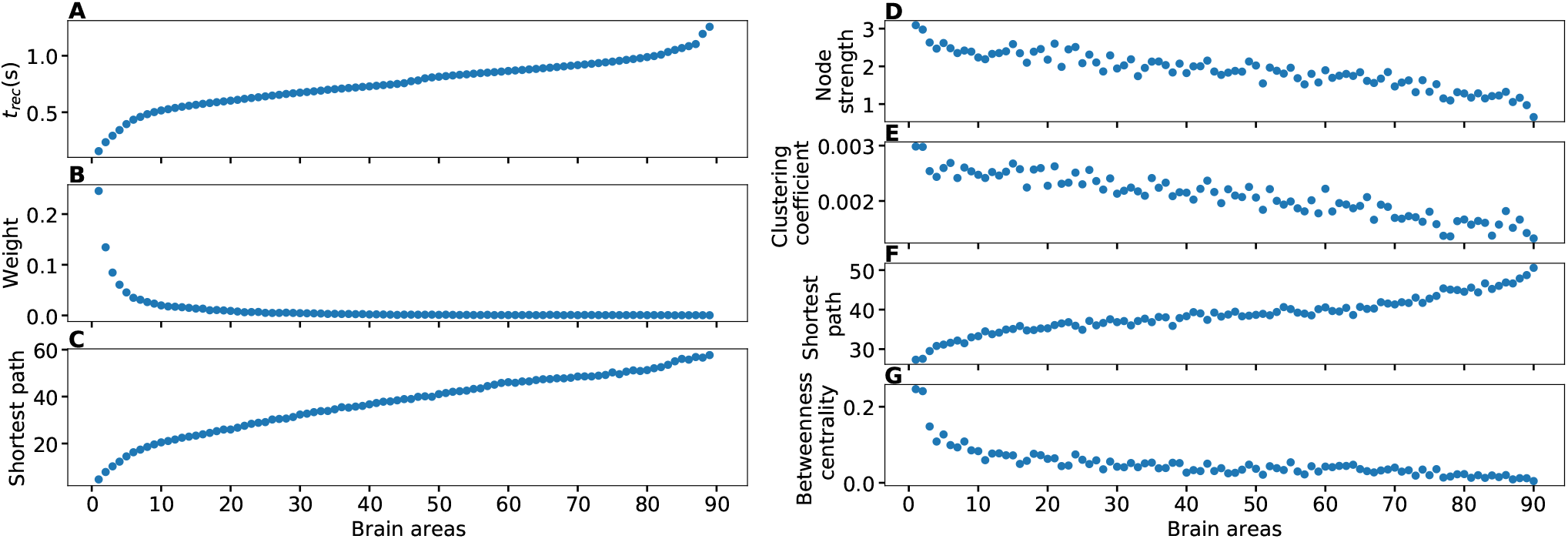
A) Recruitment times reported in descending order: Brain area 1 is the brain area which is recruited first and brain area 90 is the last recruited brain area. B) Connection weights between the recruited brain area and the EZ, ordered according to their recruitment time, thus following the indexing of panel A). C) Shortest path between the recruited area and the EZ, ordered according to their recruitment time. D) Connection weights between the recruited brain area and all the nodes except EZ, ordered according to their recruitment time. E) Clustering coefficient between the recruited brain area and all the nodes except EZ, ordered according to their recruitment time. F) Shortest path between the recruited area and all the other nodes except EZ, ordered according to their recruitment time. G) Betweenness centrality between the recruited brain area and all the nodes except EZ, ordered according to their recruitment time. The excitability 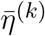 is set to the subject-specific threshold 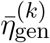, according to Fig. 3 B) for each subject separately. Data are averaged over all subjects and all the stimulated areas. Parameters: *N* = 90, Δ = 1, *σ* = 1, *I_S_* = 10, *t_I_* = 0.4 s as in Fig. 3.

On average, the first recruited brain area (labelled as 1) is connected to the EZ with a weight equal to 0.25 (1/4 of the maximum possible weight) and it is characterized by an average shortest path length to the EZ of less than 4.7. Moreover the area is recruited within an average time of less than 156 ms (calculated after the onset of the external perturbation current). However the first recruited area has, not only the strongest weight and the shortest path to the EZ but it also has, in general, the largest node strength, largest clustering coefficient, shortest average path length and largest betweenness centrality. Clearly, the seizure-like event spreads rapidly along the brain areas with strongest connection weights outgoing from the EZ; to the stronger weights are associated the shortest paths from the EZ. Overall, a region well connected is a region well recruited; this is related to the log-normal distribution of the weights (see Supplementary Fig. 2): few connections per node have a strong weight, thus allowing for fast recruitment. Note that the results for one exemplary subject and just one perturbed brain area per time (i.e. not averaged over all the brain areas and over all subjects) are comparable, even though the corresponding relationships are characterized by more variability (data not shown).

If we vary the amplitude *I*_S_ of the perturbation current, the recruitment time will vary accordingly, decreasing for increasing *I_S_*. In particular, in Fig. 8 we show an exemplary case, obtained from the stimulation of one brain area (45), for a specific subject (results are similar for other trials). Irrespectively of the recruitment order, the time needed by the first ten recruited brain areas to pass from the LA to the HA state decreases slightly for increasing amplitude. However, this decrease reaches a saturation at a current value *I*_S_ ≈ 40 already. The order of recruitment varies little: we observe some exchanges between the 4*-th* and 5*-th* and between the 9*-th* and 10*-th* recruited areas. For example, for an amplitude *I_S_* = 15, the 9*-th* recruited area (dark blue circles) gets recruited earlier than the 10-*th* area (pink dots) while, for very strong currents (*I_S_* = 100), the 9*-th* area gets recruited latest. On the other hand we do not observe a significant change in the recruitment time and order if we increase the duration *t*_I_ of the stimulation (see Supplementary Fig. 3).

**Figure 8.**
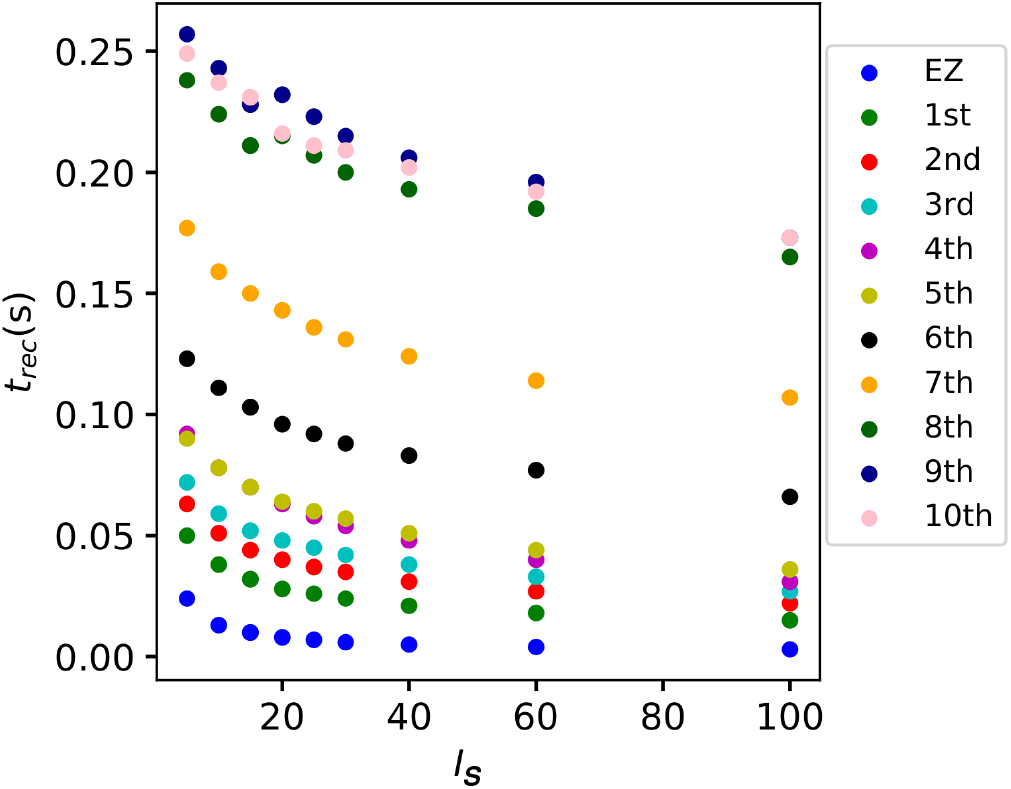
Recruitment times of the first 10 recruited areas as a function of the input current *I_S_*. The strength of the input current is varied between 0 and 100 on the x-axis. The order of the recruitment is color coded for each current strength and it changes slightly with different current strengths. Parameters: *N* = 90, Δ = 1, *σ* = 1, *t_I_* = 0.4 s, 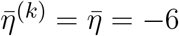, stimulation site: brain area *k* = 45 of subject 0.

### 3.2 Epileptic Patients

#### 3.2.1 Phase and Bifurcation Diagrams

In this section the structural connectivity matrices of epileptic patients are employed and an analysis, analogous to the one in Sec. 3.1.1, is provided. We present the phase and bifurcation diagrams for the multipopulation neural mass model, here employing the structural connectivity matrices of epileptic patients. As detailed before, the bifurcation diagrams shown in Fig. 9 A1)-A3), for different *σ* values, are obtained by performing an adiabatic scan along 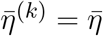, following the up- and down-sweep protocols.

**Figure 9.**
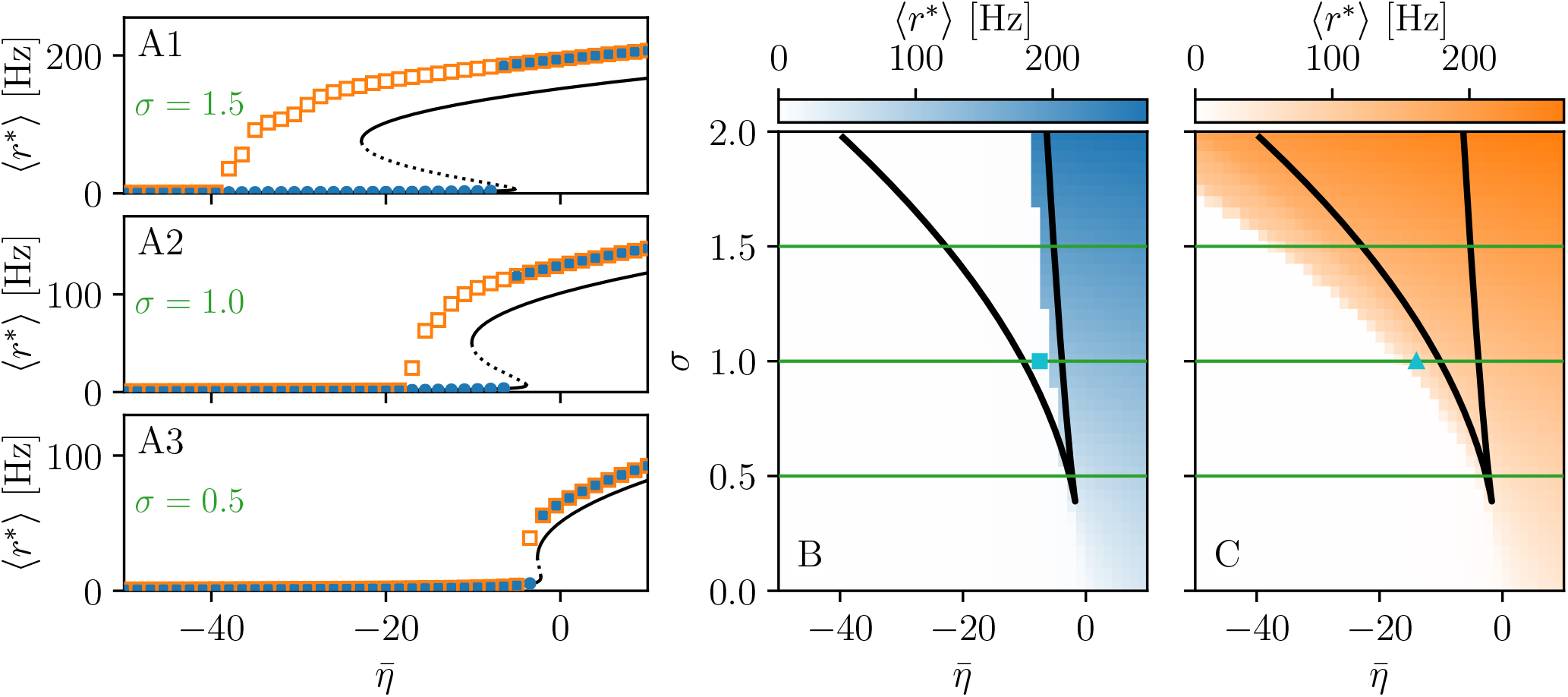
Phase and bifurcation diagrams for patient FB. A1-A3 Equilibrium firing rates 〈*r*\*〉 vs. 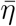 for the up-sweep (blue dots) and down-sweep (orange squares). For each 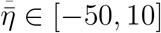 in steps of 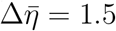 the system is initialized using the final state of the previous run and evolves for 2 s after which the average network firing rate in the equilibrium state is determined. Different panels correspond to different *σ* values: *σ* = 1.5 (A1), *σ* = 1 (A2), *σ* = 0.5 (A3). The solid (dashed) black line corresponds to the stable (unstable) equilibria in the single-node case. Maps of regimes as a function of *σ* and 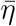 showing the network average 〈*r*\*〉 color coded for up- (B) and down-sweep (C), obtained by following the same procedure as in A1-A3 for *σ* ∈ [0, 2] in steps of Δ*σ* = 0.05. The black line indicates the single-node map of regimes like in (Montbrió et al., 2015). In panels B-C the cyan square and triangle mark 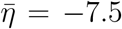 −14 respectively. Parameter values: *N*_pop_ = 88, *τ*_m_ = 20 ms, Δ = 1, *J_kk_* = 20, 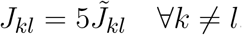.

As for the healthy subjects, the transition is hysteretic with 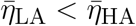. However in this case, the width of the hysteretic transition is bigger, especially for larger *σ* values. This increased width can be translated in terms of the extension of the multistability region in the phase diagram (see Fig. 9 B, C), which turns out to be slightly larger than before. The increase in size mainly occurs due to a shift of 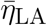, i.e. of the left boundary of the multistability regime. In this region, the transition from HA to LA, following the down-sweep, is more smooth and elongates towards smaller 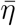 values. This implies that, in this transition region, more single population HA states exist for epileptic patients than for healthy subjects. In other words, brain areas of epileptic subjects are more prone to recruitment^2^.

While the phase diagram is obtained in the absence of time-varying input, we investigate the response of the multipopulation system to transient stimulation in the following. As for the healthy subjects, a single population is excited by injecting a step current *I*_S_(*t*) of amplitude *I*_S_ = 10 and duration *t*_I_ = 0.4 s. Initially (*t <* 0), the system is in a multistable regime, starting in the low-activity network state. For small 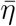 values (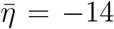, identified by the triangle in Fig. 9C), when a single node is stimulated, the system reacts analogously to the healthy subject case: During the stimulation only one stable network state exists, i.e. a focus equilibrium with a LA-HA configuration for which only the stimulated node is in HA. This focus is approached via damped oscillations (0 s *< t <* 0.4 s). When the stimulation is removed, the network maintains the LA-HA configuration, but approaches the new location of the focus again via damped oscillations. As a result, the stimulated node has large firing activity, while the remaining network is in a LA regime. For higher excitability values (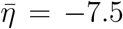, identified by the square in Fig. 9B), the perturbation of a single node gives rise to a cascade of recruitments, where other brain areas, initially not perturbed, reach the HA regime by showing damped oscillations (panels B2, B3). With respect to the recruitment features shown in Fig. 2, we observe here a faster emergence of the generalized seizure-like event: once a brain area is stimulated, the others react, in substantial number, quite immediately.

Looking at the spectrograms, the transition of the stimulated population to the HA regime is characterized by a transient activity at low frequency (*<* 20 Hz) and a sustained activity in the *γ* band (50-180 Hz), observable throughout the duration of the stimulus, as shown in panel A5, where the spectrogram for the single stimulated population is reported. Regarding the spectrogram of the mean membrane potentials averaged over the network population (panel A4), it turns out that the low frequency activity in the *δ*, *θ* bands is present, while the activity at high frequency simply reflects the activity of the stimulated area. In the case of large recruitment events, at larger excitability values, it is possible to observe *γ* activity at higher frequencies (see panels B4-B5), which is enhanced with respect to the situation where an asymptomatic seizure-like event is present. Moreover, comparing the spectrograms in Fig. 10 and those reported in Fig. 2, we see that the activity takes place at higher frequency ranges when considering structural connectivity matrices of epileptic patients and the activity is mainly concentrated in the EZ. The last statement may be qualified, however, by recent studies proposing high frequency oscillations (80-500 Hz) recorded not only at seizure onset but also between seizures (the interictal period), as a putative new marker of the epileptogenic focus Jacobs et al. (2012). More specifically fast cortical ripples superimposed to interictal epileptiform discharges were correlated with the seizure onset zone and primary propagation area in neocortical epilepsy Khadjevand et al. (2017). Neocortical ripples were also found to be more specifically confined to the seizure onset and propagation regions, and thus a better marker compared to interictal epileptiform discharges alone Wang et al. (2013). High frequency oscillations, as shown in Fig. 10 B4, B5, are much more frequent in the seizure-like onset zone than outside, where they are often totally absent. The rather empty spectrograms of mean membrane potentials for patient FB are a result of a rather rapid recruitment of a majority of nodes, thus giving rise to a strong signal change, immediately upon recruitment, which suppresses the rest of the signal in the spectrogram. At the same time the damped oscillations are all compressed within a narrow time window, and not very elongated in time, as it happens for healthy subjects (see Fig. 2). In other words, if the generalized seizure-like event is rapid, all the signals overlap, and this is especially clear looking at the strong low frequency bands. A fast generalized seizure-like event, in absence of high frequency oscillations outside the EZ, can be obtained for healthy subjects only increasing the excitability parameter: for higher 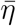 values, the recruitment is more sudden, as shown in the Supplementary Fig. 21.

**Figure 10.**
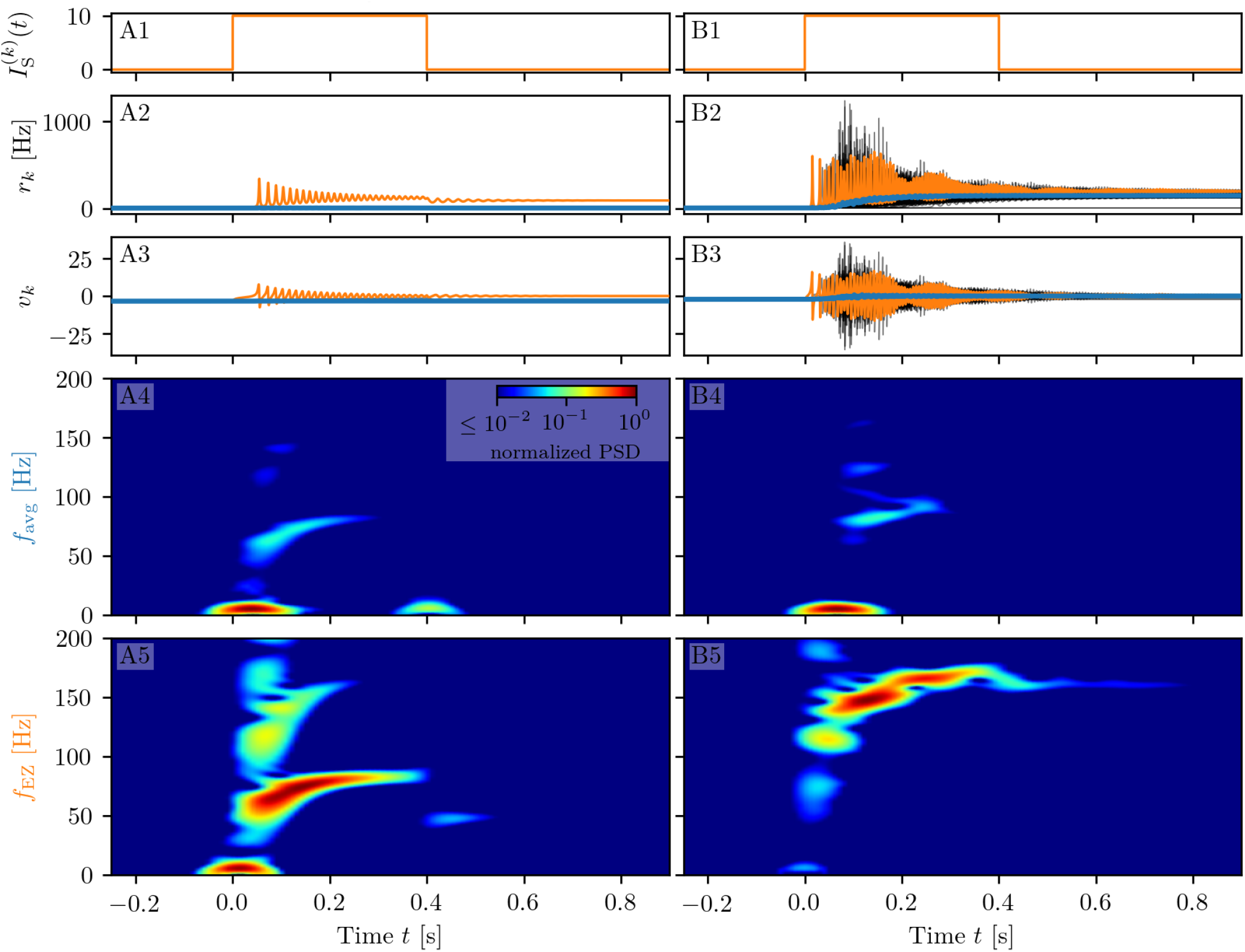
Spectrograms of mean membrane potentials for patient FB. (A1-B1) Stimulation current 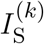, (A2-B2) population firing rates *r_k_* and (A3-B3) mean membrane potentials *v_k_* for the EZ (orange) and other populations (black). The blue curves show the network average firing rate and membrane potential. (A4-B4) Spectrogram of the network average membrane potential and (A5-B5) of the *v_k_* of the EZ. Column A shows an asymptomatic seizure-like event for 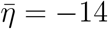, column B a generalized seizure-like event for 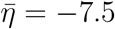. In both cases the EZ node 46 is stimulated. Parameter values: *N*_pop_ = 88, *τ*_m_ = 20 ms, Δ = 1, *σ* = 1.25, *J_kk_* = 20, 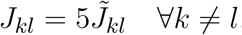.

#### 3.2.2 Temporal Recruitment of Clinically and SEEG Predicted Propagation Zones

In the following we test the clinical predictions for epileptic patients, by choosing the EZs, identified by clinical doctors via presurgical invasive evaluation, as perturbation sites. We investigate the recruitment times of different brain areas following such a perturbation and compare the order of recruitment with the experimental data given for each individual subject. A general overview on the recruitment times of all brain areas, for all patients, is shown in Fig. 11. As perturbation sites, the clinical EZs, are used for all patients. The perturbation step current (*I*_S_ = 10, *t*_I_ = 0.4 s) is applied, to each perturbation site, in correspondence with the dashed vertical black line. The parameters are identical for almost all patients and are chosen such that at least 90% of the brain areas are recruited while still allowing multistability among various LA-HA configurations, including the network LA state. For each patient (identified via the initials on the y-axis), the recruitment time of each brain area is reported: The grey dots represent the time values for each brain area. Superimposed on the grey dots are red × and blue + marks that identify the brain areas belonging to the PZ, according to the non invasive (PZ_Clin_) or invasive (PZ_SEEG_) presurgical evaluation, respectively. The recruitment time averaged over all brain areas is identified, for each patient, by a green vertical line, while the boxes contain the second and third quartile of the distribution, and the whiskers have 1.5 the length of the InterQuartile Range (IQR) from the upper or lower quartiles. Remarkably, the propagation zones PZ_Clin_ and PZ_SEEG_ turn out to be among the first recruited brain areas for all patients in the numerical experiments. However the temporal dynamics vary for all patients, with GC and AC having late recruitments. Looking at the set of the first ten recruited brain areas for each patient (reported in detail in the Tables 5-7 in the Supplementary Material), we notice that most of the areas, identified by clinicians as belonging to the PZ, are actually within this set: For patients CV, ET, FB, IL, SF all the areas belonging to PZ_Clin_ are among the first ten recruited areas, while the same holds true for patients CJ, CM, FB if we consider the areas identified by the stereotactic EEG analysis as belonging to the propagation zone (PZ_SEEG_). In general a large number of the first ten recruited areas, as revealed by our simulations, coincides with the areas that are supposed to be crucial in the seizure spreading according to the medical doctors (e.g. for patients CJ, CM, JS, PC, PG, RB). Moreover the predictabiliy of the model is higher if we compare our results with the predictions PZ_Clin_. Finally, the brain areas belonging to the predicted propagation zones, are in general recruited before the median recruitment time.

**Figure 11.**
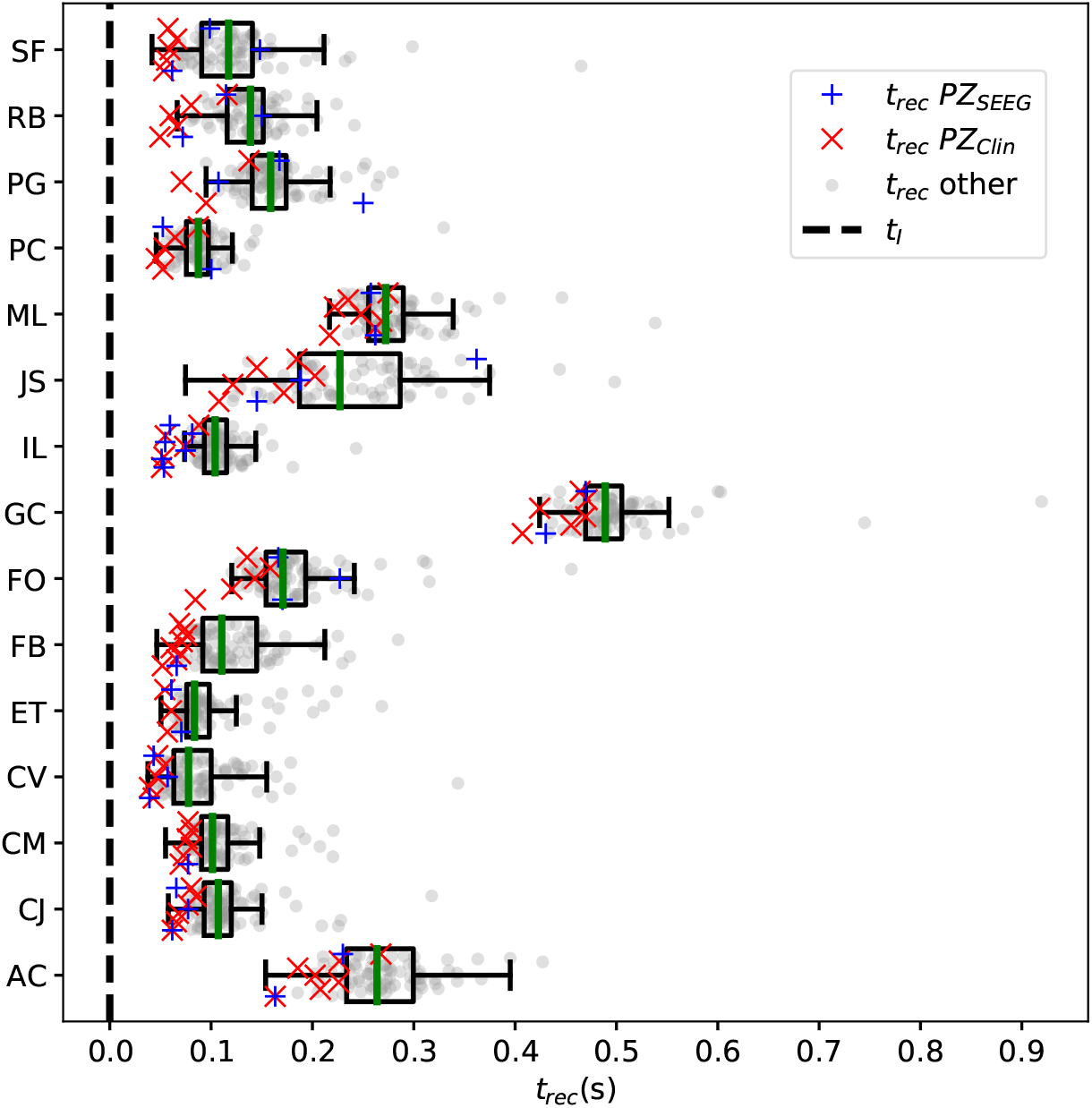
Recruitment times of all brain areas for the cohort of epileptic patients: The recruitment time, reported on the x-axis, identifies the time needed by a brain area to jump to the HA regime after the application of the perturbation current. The boxplots consist of the recruitment times of all brain areas for each patient. Patients are identified according to their initials on the y-axis. The median is represented as a green vertical line while the boxes contain the second and third quartile of the distribution. The whiskers are chosen with maximum length 1.5 × IQR and show the most extreme observed values that are within 1.5 × IQR from the upper or lower quartiles. The grey dots represent the recruitment times for each brain area. The red × shows the recruitment of a brain area clinically predicted to be part of the propagation zone PZ_Clin_. The blue *+* represents the recruitment of a brain area which is part of the propagation zone according to the SEEG measurements PZ_SEEG_. Parameters: *N*_pop_ = 88, Δ = 1, *σ* = 1.25, *I_S_* = 10, *t_I_* = 0.4 s, 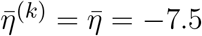 (except for patients AC 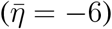 and ML 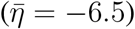).

**Table 5.** List of the first 10 recruited brain areas for each patient. The column “Type” indicates whether the recruited area belongs or not to the PZ estimated via presurgical invasive (PZ_*SEEG*_) or non-invasive (PZ_*Clin*_) evaluation.

| Patient | Recruitment order | Recruitment time (s) | Type | Region |
| --- | --- | --- | --- | --- |
| AC | 0 | 0.0 | EZ | rh-LOFC |
| AC | 1 | 0.0005 | EZ | rh-TmP |
| AC | 2 | 0.123 | PZ<sub>SEEG</sub>, PZ<sub>Clin</sub> | rh-RMFG |
| AC | 3 | 0.1324 | PZ<sub>Clin</sub> | rh-Pu |
| AC | 4 | 0.1546 | other | rh-SFG |
| AC | 5 | 0.1587 | PZ<sub>Clin</sub> | rh-Ins |
| AC | 6 | 0.1718 | other | rh-Pal |
| AC | 7 | 0.1747 | other | rh-PrG |
| AC | 8 | 0.175 | other | rh-MOFC |
| AC | 9 | 0.1769 | other | rh-Cd |
| AC | 10 | 0.1797 | other | lh-SFG |
| AC | 11 | 0.1801 | other | rh-PoG |
| CJ | 0 | 0.0 | EZ | lh-LOCC |
| CJ | 1 | 0.0255 | PZ<sub>SEEG</sub>, PZ<sub>Clin</sub> | lh-FuG |
| CJ | 2 | 0.0294 | PZ<sub>SEEG</sub>, PZ<sub>Clin</sub> | lh-SPC |
| CJ | 3 | 0.0333 | PZ<sub>Clin</sub> | lh-ITG |
| CJ | 4 | 0.0354 | PZ<sub>SEEG</sub>, PZ<sub>Clin</sub> | lh-IPC |
| CJ | 5 | 0.045 | other | lh-MTG |
| CJ | 6 | 0.046 | other | lh-SMG |
| CJ | 7 | 0.0466 | other | lh-PCunC |
| CJ | 8 | 0.0479 | PZ<sub>Clin</sub> | lh-LgG |
| CJ | 9 | 0.0482 | other | lh-PoG |
| CJ | 10 | 0.0521 | other | lh-PrG |
| CM | 0 | 0.0 | EZ | lh-Ins |
| CM | 1 | 0.0224 | PZ<sub>Clin</sub> | lh-Pu |
| CM | 2 | 0.0369 | other | lh-SFG |
| CM | 3 | 0.0394 | PZ<sub>Clin</sub> | lh-LOFC |
| CM | 4 | 0.04 | PZ<sub>Clin</sub> | lh-PrG |
| CM | 5 | 0.0438 | PZ<sub>SEEG</sub>, PZ<sub>Clin</sub> | lh-PoG |
| CM | 6 | 0.0446 | other | lh-RMFG |
| CM | 7 | 0.0451 | other | lh-Th |
| CM | 8 | 0.0453 | other | lh-CMFG |
| CM | 9 | 0.0459 | other | rh-SFG |
| CM | 10 | 0.0465 | PZ<sub>Clin</sub> | lh-Pop |
| CV | 0 | 0.0 | EZ | lh-SFG |
| CV | 1 | 0.0016 | EZ | lh-CMFG |
| CV | 2 | 0.0026 | EZ | lh-PCG |
| CV | 3 | 0.007 | PZ<sub>SEEG</sub>, PZ<sub>Clin</sub> | lh-PrG |
| CV | 4 | 0.0085 | PZ<sub>Clin</sub> | lh-RMFG |
| CV | 5 | 0.0122 | PZ<sub>SEEG</sub> | lh-PoG |
| CV | 6 | 0.0126 | PZ<sub>Clin</sub> | rh-SFG |
| CV | 7 | 0.0155 | PZ<sub>Clin</sub> | lh-PaC |
| CV | 8 | 0.0168 | other | lh-Pop |
| CV | 9 | 0.0168 | other | lh-Pu |
| CV | 10 | 0.0175 | other | lh-Th |
| CV | 11 | 0.0211 | other | lh-SMG |
| CV | 12 | 0.0212 | PZ<sub>Clin</sub> | lh-CACC |
| ET | 0 | 0.0 | EZ | lh-PCunC |
| ET | 1 | 0.0006 | EZ | lh-PCG |
| ET | 2 | 0.0181 | PZ<sub>Clin</sub> | lh-SPC |
| ET | 3 | 0.0219 | PZ<sub>Clin</sub> | lh-ICC |
| ET | 4 | 0.0244 | PZ<sub>SEEG</sub>, PZ<sub>Clin</sub> | lh-IPC |
| ET | 5 | 0.0284 | other | lh-SFG |
| ET | 6 | 0.0318 | other | lh-LOCC |
| ET | 7 | 0.0345 | other | rh-SFG |
| ET | 8 | 0.0363 | other | lh-Cun |
| ET | 9 | 0.0364 | other | lh-SMG |
| ET | 10 | 0.0366 | other | lh-Th |
| ET | 11 | 0.0374 | other | lh-PrG |

**Table 6.**
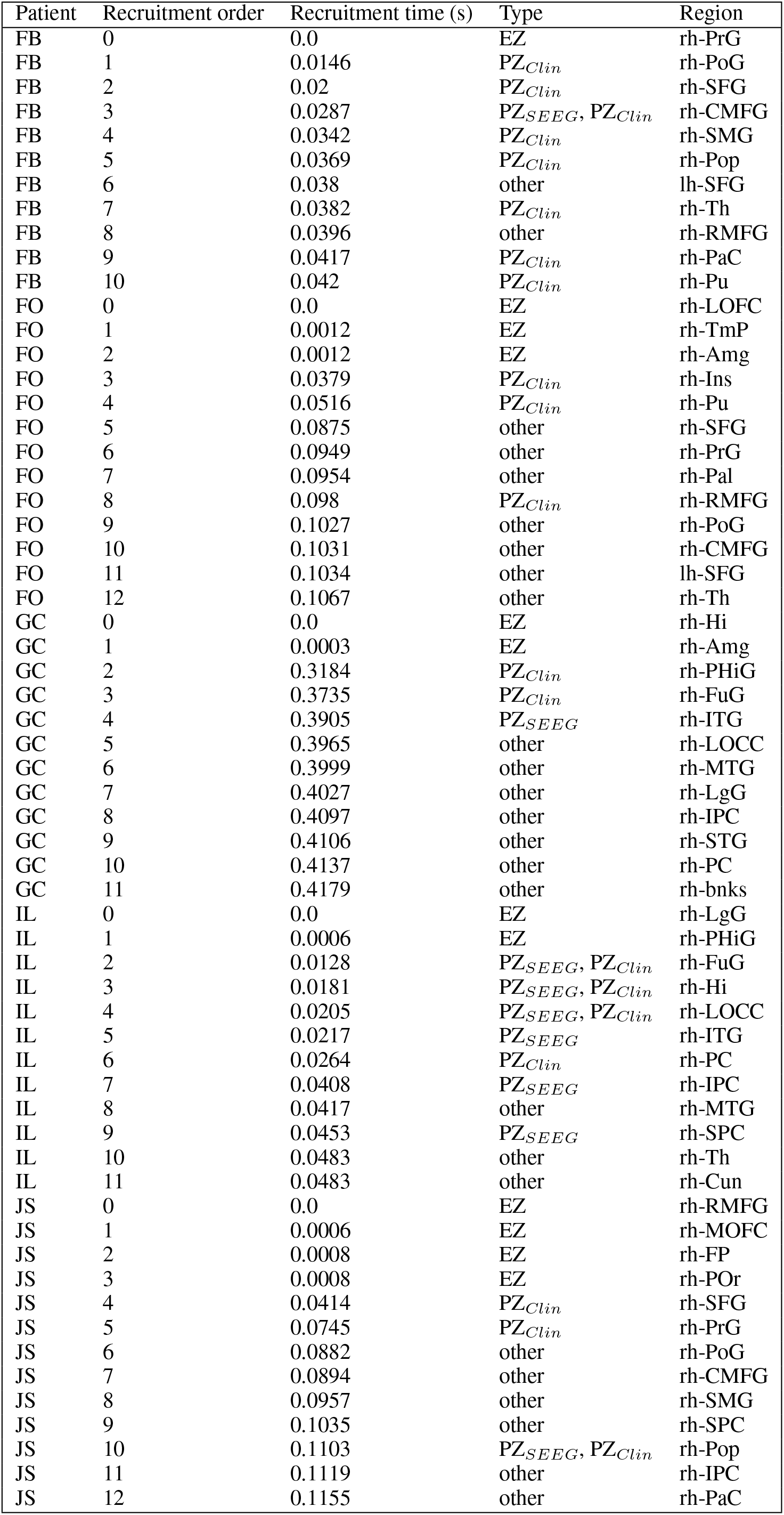
Continued from Table 5.

**Table 7.**
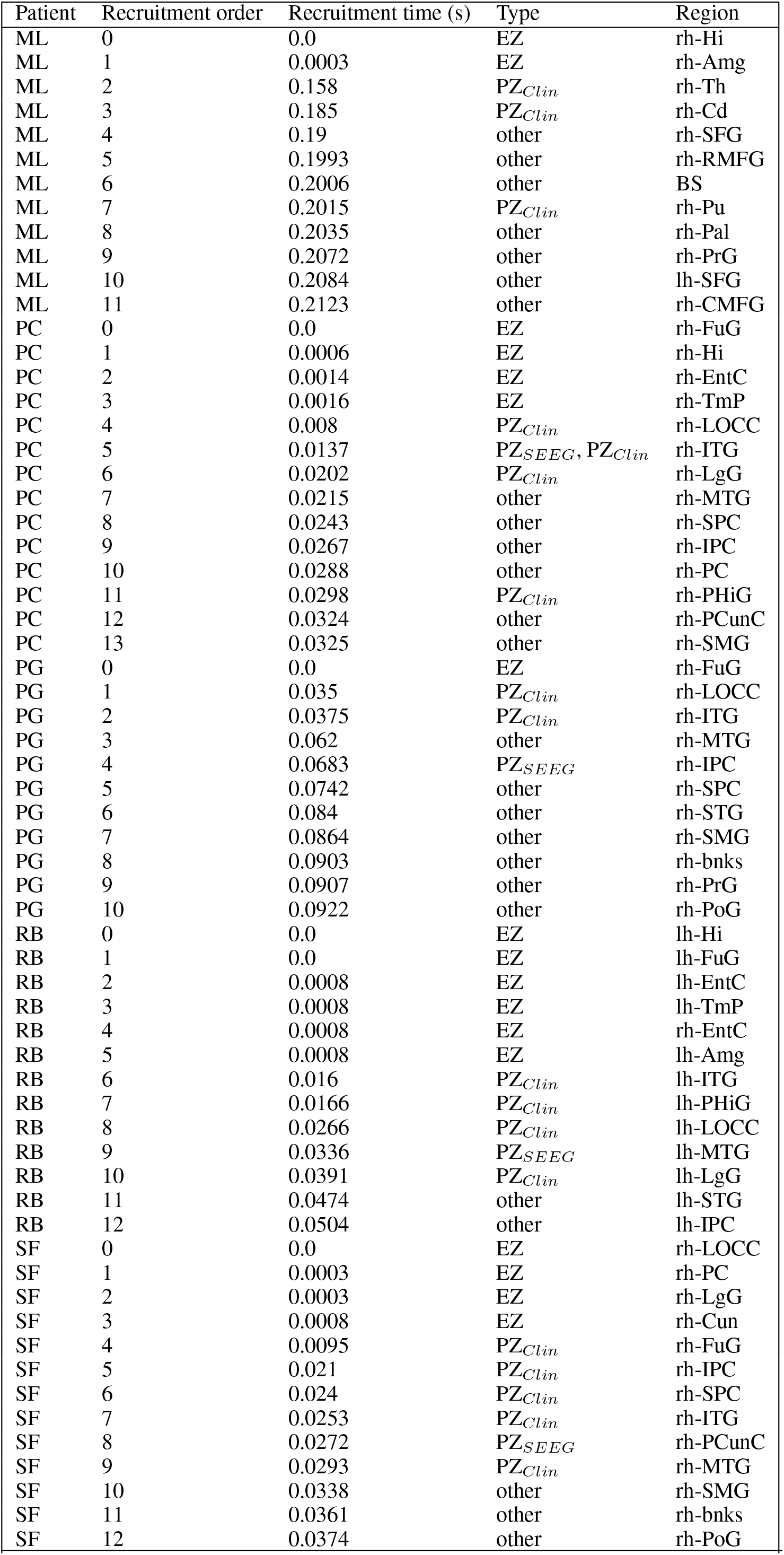
Continued from Tables 5, 6.

To evaluate the dependence of the shown results on the chosen parameters, with the idea in mind of going towards a more biologically realistic framework, we have repeated the previous numerical experiment by employing a random Gaussian distribution of the excitability parameter 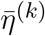(see Fig. 12). The distribution is centred at 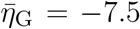 with standard deviation 0.1 for all patients except AC and ML. For the latter patients we shifted the center towards larger values, in order to get a sufficient number of recruitments when the EZ is stimulated. In all cases the results are averaged over 10 different random realizations of the distribution. More details on the impact of different realizations of 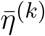 are given, for one exemplary patient, in the Supplementary Fig. 22. For larger standard deviations than the one employed, a too large fraction of the populations would not be able to exhibit bistability between LA and HA, highlighting the system sensitivity to small parameter changes. However, for the chosen distribution, the results are comparable with the ones obtained with identical 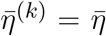, shown in Fig. 11. For patients CJ, CM, CV, ET, FB, IL the predicted propagation zones are always the first ones to be recruited. Moreover most of the areas are usually recruited in the first half of the recruitment process, rapidly increasing in number, once the areas in the propagation zones have been recruited (thus giving rise to a peak in the histogram). As a general remark, in view of the distributed nature of the excitabilities, recruitments at later times, with respect to the former case with homogeneous 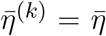, may now take place.

**Figure 12.**
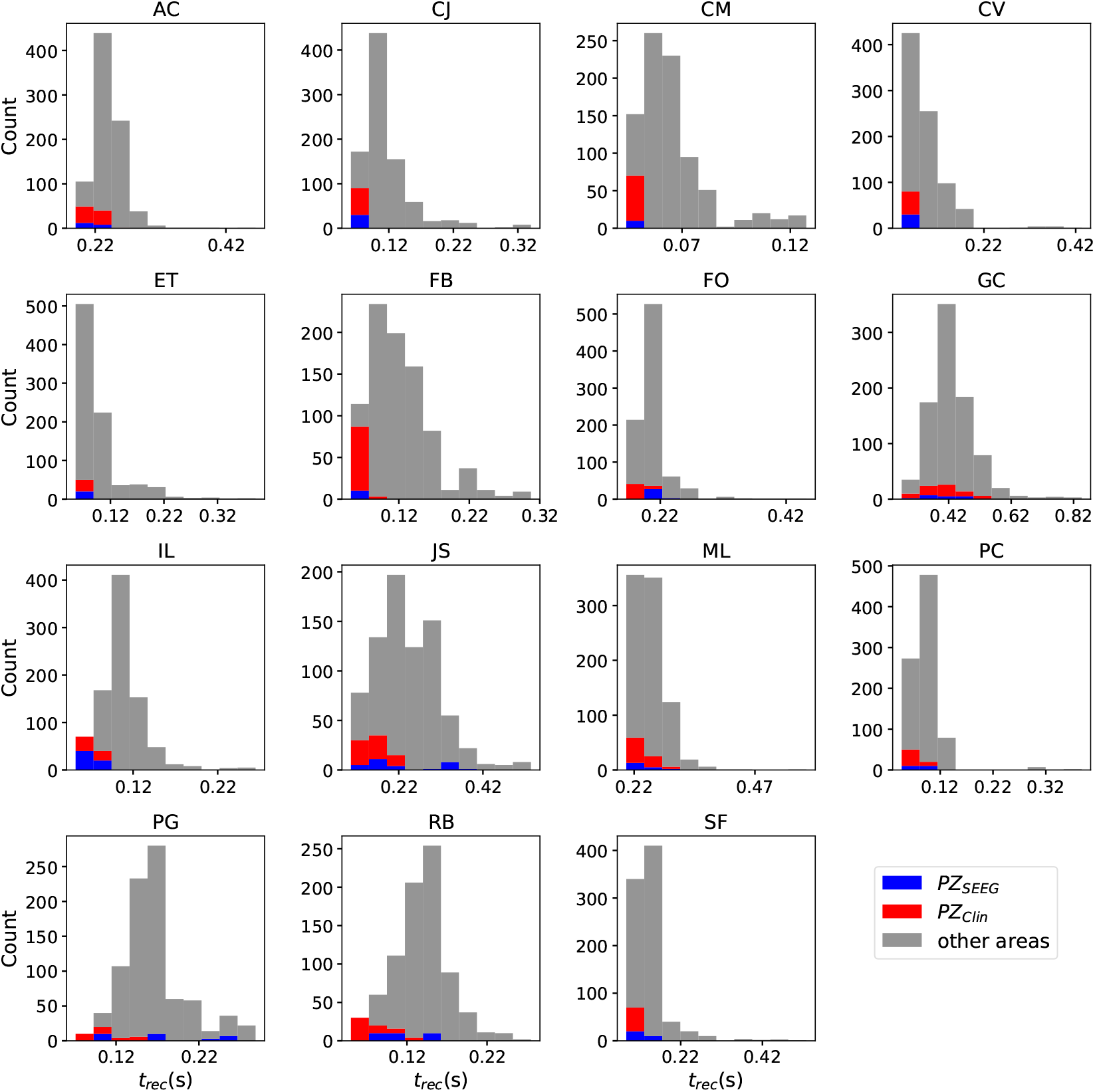
Histograms of recruitment times for all epileptic patients. For each patient (identified by his/her initials), the recruitment times of all the brain areas are collected, once the EZ is stimulated. The EZ is chosen according to the presurgical evaluation (see Table 4 of the Supplementary Material) and vary from one patient to the other. Parameters as in Fig. 11 except for 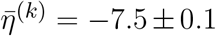 (for AC 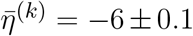, for ML 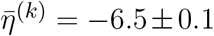. Results are averaged over 10 repetitions of different random Gaussian distributions.

**Table 3.** Clinical characteristics of the patients. N, normal; L, left; R, right; Th, thermocoagulation; Gk, Gamma knife; Sr, surgical resection; NO, not operated; PVH, periventricular nodular heterotopia; FCD, focal cortical dysplasia; SPC, superior parietal cortex; F, Frontal; NA, not available.

| Patient | Gender | Epilepsy duration (years) | Age at seizure onset (years) | Epilepsy type | Surgical procedure | Surgical outcome | MRI | Histopathology | Side |
| --- | --- | --- | --- | --- | --- | --- | --- | --- | --- |
| AC | F | 14 | 8 | Temporo-frontal | Sr | III | Anterior temporal necrosis | Gliosis | R |
| CJ | F | 14 | 9 | Occipital | Sr | III | N | FCD type 1 | L |
| CM | M | 35 | 7 | Insular | GK | I | N | NA | L |
| CV | F | 18 | 5 | SMA | Sr | I | N | FDC type 2 | L |
| ET | F | 23 | 7 | Parietal | Sr | I | FCD SPC | FCD type 2 | L |
| FB | F | 16 | 7 | Premotor | Th | II | N | NA | R |
| FO | M | 45 | 11 | Temporo-frontal | Sr | I | FCD F | FCD type 2 | R |
| GC | M | 5 | 28 | Temporal | Sr | III | Temporopolar hypersignal | FCD type 1 | R |
| IL | F | 18 | 20 | Occipital | N | NO | N | NA | R |
| JS | M | 11 | 18 | Frontal | Sr | I | Frontal necrosis (post-trauma) | Gliosis | R |
| ML | F | 10 | 17 | Temporal | Gk | II | Hippocampal sclerosis | NA | R |
| PC | M | 15 | 14 | Temporal | N | NO | N | NA | R |
| PG | M | 29 | 7 | Temporal | Sr | I | Cavernoma | Cavernoma | R |
| RB | M | 28 | 35 | Temporal | Sr | III | N | Gliosis | L |
| SF | F | 24 | 4 | Occipital | N | NO | PVH | NA | R |

**Table 4.**
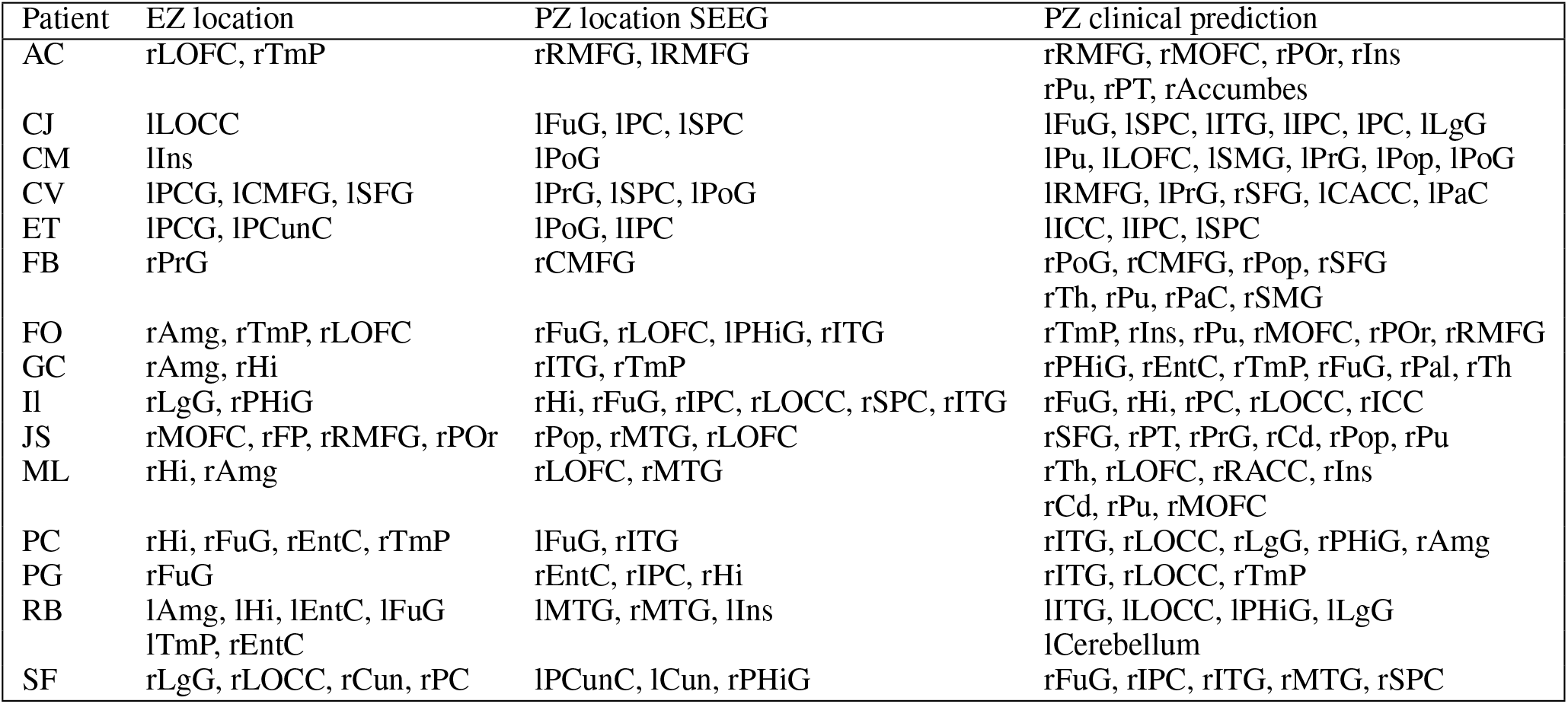
Results of Propagation zone prediction for each patient. Abbreviations are given in Supplementary Table 2.

For patients with many nodes in the EZ, the recruitment process may result to be more complex, as it happens for patients RB and JS, for which the histograms are less narrow, but instead widely distributed. However this cannot be taken as a general rule, since comparable histograms are obtained for patients PG (one node in the EZ) and GC (two nodes in the EZ), while for SF and PC (with both four nodes in the EZ) the histograms result to be very narrow, thus implying a fast recruitment process of most of the brain areas. The differences among the histograms can be partially justified by the facts that patients have specific connectomes with individual characteristics and by the analysis that we have proposed by choosing similar 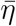 values for all the patients. In this way we have preferred to have a general look on the multiple self-emergent dynamics in a group of patients, instead of fine-tuning the excitability parameter in order to obtain similar collective behaviors. What we observe here is strongly related to what we have presented in Fig. 10 and in the Supplementary Fig. 21: the recruitment speed depends on the excitability parameter and on the individual network structure. Faster recruitment events may be obtained for different subjects by increasing the excitability value.

#### 3.2.3 Relationship Between DTI Network Structure and Temporal Seizure Recruitment

In order to understand the mechanism underlying the recruitment events, we evaluate the relationship between the network structure, in terms of topological measures, and the recruitment times of the first 10 recruited brain areas. For simplicity, we consider patients with only one brain area in the EZ and we report, in Fig. 13, the EZ (purple dot) and the first 10 recruited areas in a graph representation. The first recruited areas are ordered according to their recruitment times in clockwise order. Moreover we indicate in blue the areas belonging to the PZ, as identified according to the presurgical invasive evaluation (PZ_SEEG_). Black lines identify the weighted connections between all areas and their thickness is proportional to their weight. The sizes of the circles representing each brain area are proportional to their inverse recruitment time (A1-D1), to their weight connecting each area to the EZ (A2-D2), and to their inverse shortest path length between each node and the EZ (A3-D3), while the size of the purple EZ circle remains fixed.

**Figure 13.**
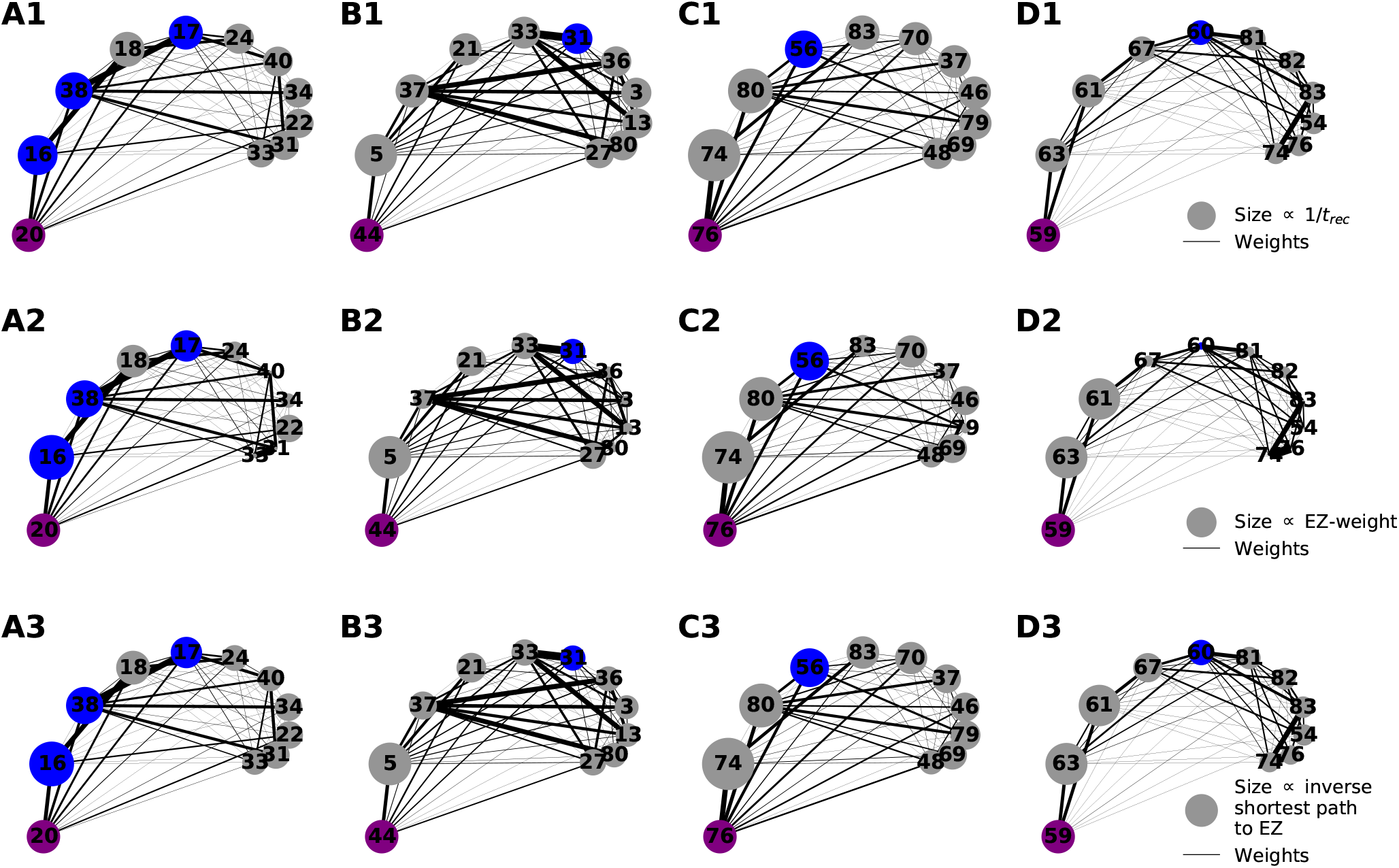
Graph plot of the first 10 recruited areas, ordered clockwise according to their recruitment times. Node circle size corresponds to the inverse recruitment time (A1-D1), to the connection strength to the EZ (A2-D2) and to the inverse shortest path length to the EZ (A3-D3). The purple dot identifies the EZ and its size remains fixed. Blue dots distinguish a recruited area to belong to the PZ_SEEG_, i.e. the PZ identified according to the presurgical invasive evaluation. Results are obtained for patients CJ (panels A1-A3), CM (panels B1-B3), FB (panels C1-C3), PG (panels D1-D3). Parameters as in Fig. 11.

Since in (A1-D1) the node size is proportional to the inverse recruitment time, large circles indicate early recruitment while small circles indicate late recruitments: Hence the circles become smaller clockwise. In panels (A2-D2) the node size is proportional to the weight connecting each area to the EZ and it turns out that, for all patients, the first recruited area has the strongest connecting weight. However, after a few recruitments this does not hold true anymore. There are many examples in which areas with a strong weight to the EZ (see e.g. area 46 or 48 for patient FB) are recruited much later than areas with very small weights (e.g. area 83 for FB). The seizure-like event propagates as a chain reaction and, therefore, the strongest connecting weight to the EZ is only decisive for the very first recruited area. Later, strong connections to other early recruited areas play a decisive role, as it is the case for area 83 in FB which has a weak connection weight to the EZ. However, through its strong connection to area 74, its weighted shortest path length to the EZ is quite short, thus meaning that the weighted shortest path length to the EZ cannot be underestimated in order to find the recruitment ordering. Indeed, in (A3-D3) one can see the good predictability of the shortest path: the node size, proportional to the inverse shortest path length to EZ, decreases in general with later recruitment. This is expected, given the fact that the average shortest path to the EZ considers all connections in the network, not just the connections subgraph outgoing the EZ. An example of the high predictability of the shortest path is given by the node 38 in patient CJ, which has a shorter path length to the EZ than node 18. Node 38 is recruited before node 18 irrespectively of its strong connection to node 16 and a connection strength to the EZ comparable with the one of node 38.

For later recruitments, the prediction becomes even more difficult because one needs to account for the temporal order of the seizing brain areas. As shown before, the area which is first recruited, is the one with the strongest connection to the EZ. However, depending on the strength of the connection, the recruitment time changes and it increases for decreasing strength. In the case of patient CJ the recruitment of the second area is determined, more by the strength of the connections to the EZ (i.e. area 20) than by the connection to area 16, while, for the recruitments of the third and forth areas, the strong connections of node 18 to 16 and of node 17 to 38, i.e. the first and second recruited nodes, are fundamental. On the other hand, when the first recruited areas have strong connections to the EZ, as for example area 74 in patient FB, the successive recruitments are strongly influenced by the first recruited area, whose outgoing graph reveals areas that are recruited with high probability. Thus the connection to area 74 turns out to be, for the second, third, and fourth recruitment almost as important as the connection to the EZ (i.e. area 76). Finally, if we compare two late recruited areas that are characterized by the same shortest path length to the EZ, but with a path to the EZ that crosses very different nodes, we observe that the area with the path going through earlier recruited nodes is recruited earlier. The longer the seizure-like event propagates, the less important the shortest path length to the EZ becomes and the more important the path lengths to other recruited nodes become. This underlines the difficulty of predicting the seizure propagation in complex networks.

To confirm the importance of the shortest path length and the strength of the connections outgoing the EZ in determining recruitment events, we report in Fig. 14 the recruitment time values as a function of the shortest path and the connection weights for the patients with a single node as EZ (panels A, B) and for all 15 epileptic patients (panels C, D). While in panel B the recruitment time is plotted over the logarithm of the weight, in panel C (D) the values of the recruitment time, plotted as a function of the shortest path (connection weight), are ordered according to their recruitment order. In particular the order for recruitment, shortest path, and weight to EZ is ascending from small values to large values. This means that, in panel D, the areas with the strongest weights (87th, 86th, etc.) correspond to the areas that are recruited earliest (1st, 2nd, etc.). The ordering has been preferred to the specific values of the shortest path and connection weight when reporting data for all 15 patients, in order to obtain a better visualization. For patients CJ, CM, PG, FB, the recruitment time grows almost linearly with the shortest path, while it decreases for increasing weights. This analysis is confirmed in the Supplementary Fig. 23, where a regression fit is performed over the data shown in panel A, thus underlying the approximately linear relationship between the shortest path length and the recruitment time for larger *t_rec_*. The relationship is not anymore so evident when we consider different cases of EZs, that are composed of more that one area. However, in this case, it is still possible to affirm that the earliest recruitments are associated with the shortest path lengths and the strongest weights, while the nodes corresponding to PZ_SEEG_ or PZ_Clin_ that, according to our simulations, were recruited late, have very long shortest path lengths to the EZs or very small weights.

**Figure 14.**
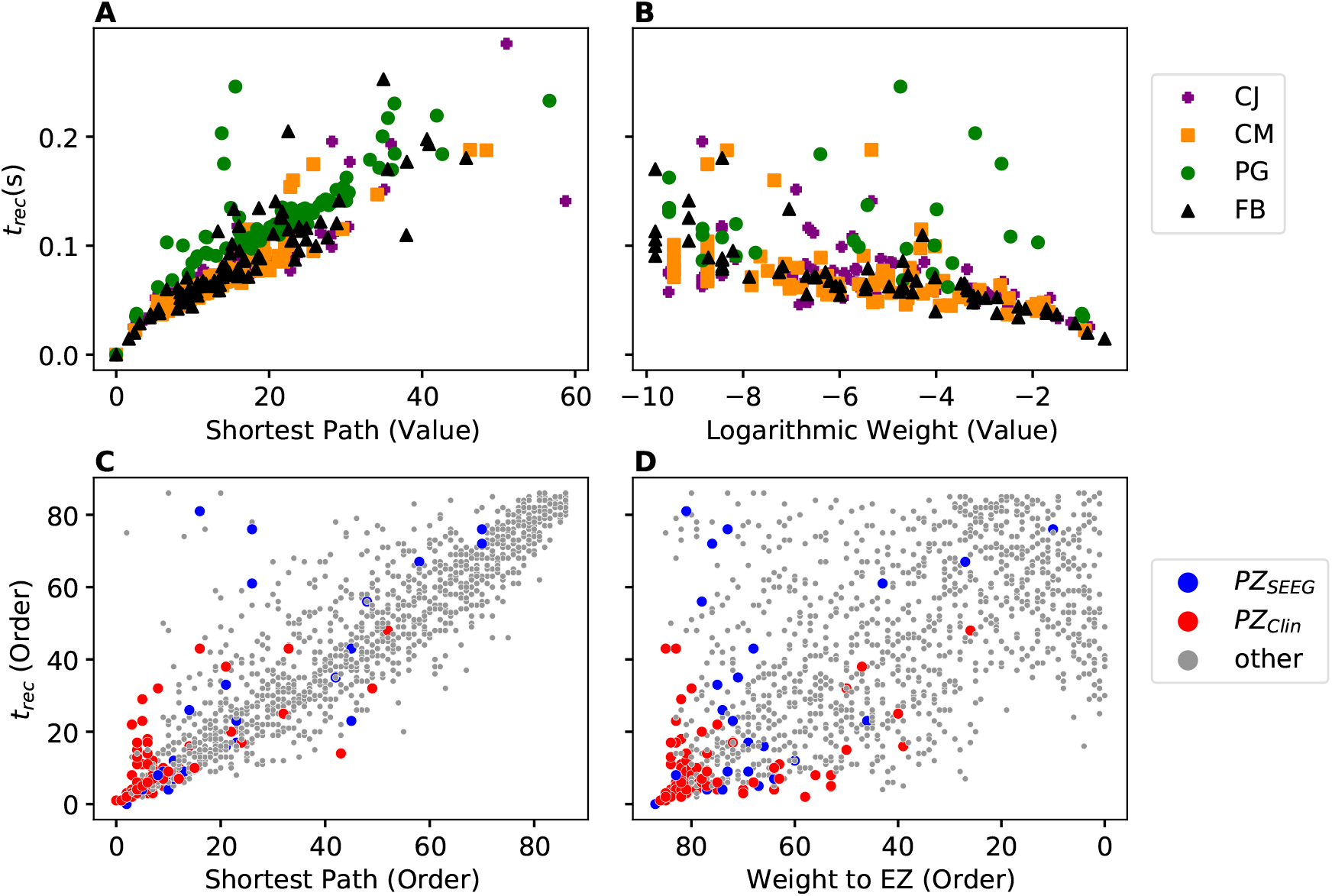
Relationship between network measure and recruitment time for four patients with one EZ: A) Shortest path to EZ; B) Logarithmic value of the weight to EZ. In A) all four EZs are shown at (0, 0) while in B) the EZs are omitted. The recruitment time is calculated, in seconds, after the perturbation current has started. In C), D) the recruitment time values are plotted according to their order, as a function of shortest path to EZ (C) and weight to EZ (D) for all 15 patients. In D) the x-axis was inverted for better comparison. Parameters as in Fig. 11.

In general the recruitment mechanism is not completely defined by the shortest path length and the connection weight, therefore it is not possible to match the pre-surgical predictions in terms of PZ_SEEG_ and PZ_Clin_ if we try to identify the nodes belonging to the PZ by calculating the first recruited nodes according to their shortest paths length or their connection weights. In particular, it turns out that the PZ_SEEG_ areas are well predicted by the investigated model if the shortest path length between the predicted PZ and the EZ is short, as shown in Fig. 15 A). However, for patients GC and JS, the recruitments of the nodes belonging to PZ_SEEG_ happen much later when compared to brain areas of other patients with a similar shortest path length. Equivalently in panel B) it is possible to observe that, for short values of the shortest path length (*<*5), there is a linear correspondence between short recruitment times and PZ_Clin_ areas that are characterized by small values of the shortest path. However the areas belonging to PZ_Clin_ are still not identifiable, in terms of topological measures, for patient GC.

**Figure 15.**
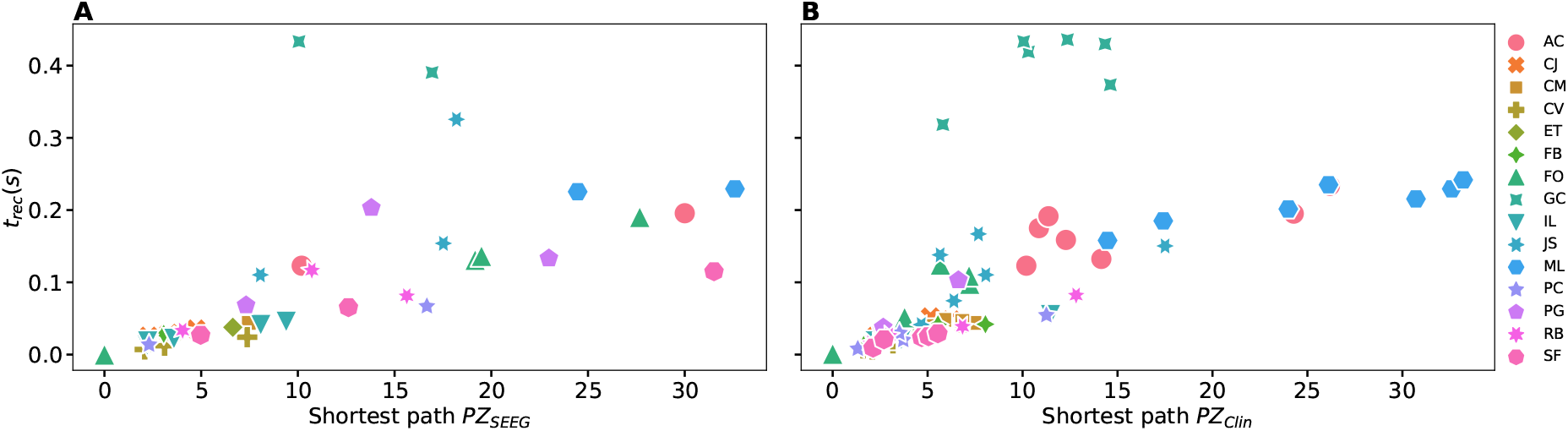
Recruitment times *t_rec_* of the areas belonging to PZ_SEEG_ (A) and PZ_Clin_ (B) as a function of the shortest path length to EZ, for all patients. Parameters as in Fig. 11.

To conclude this Section on the influence of single connectome topology in determining activity spreading and area recruitment, we elaborate the data reported in Fig. 11 by sorting, from top to bottom, the patients according to their median shortest path length, calculated on all areas with respect to the EZ. In Fig. 16 are shown the recruitment times of all brain areas for all patients. Since patients are ordered according to their median shortest path length, the brain areas of CV have, on average, the shortest paths to the EZ and the areas of AC the longest. In general, it is possible to detect a slight trend, for the overall recruitment events, to delay with longer average shortest path lengths. More in detail, JS and GC show both very long and very short recruitment times, thus confirming the results obtained in Fig. 12 for Gaussian-distributed excitabilities. The scattering of the recruitment times for these patients reflects that, on average, their recruitment times are longer with respect to the other patients. However the mean recruitment times are comparable with those of ML, AC, that show comparatively late recruitments irrespectively of the fact that are characterized by a longer median shortest path. A common characteristic that brings together patients JS, GC, ML, AC is the weak connection among the EZ and the first recruited area, that slows down the recruitment time (as already mentioned when discussing about Fig. 13), thus suggesting that is the interplay between connection strength and shortest path to determine the efficacy of seizure spreading and not the single topology measure alone.

**Figure 16.**
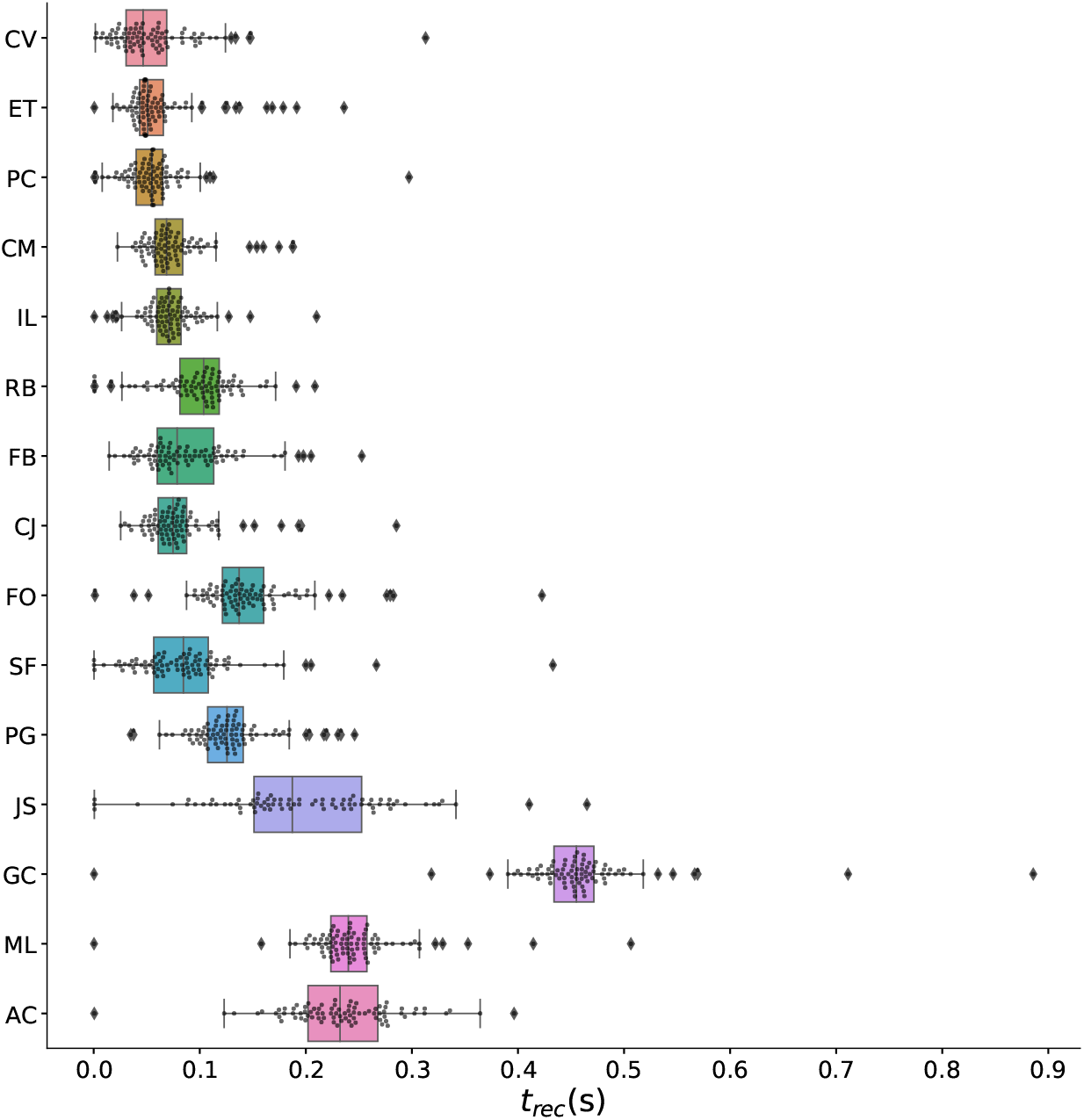
Recruitment times of all brain areas and all patients. The patients are sorted from top to bottom according to their median shortest path length, calculated by listing all the shortest path lengths of all areas to the EZ and then locating the number in the centre of that distribution. Grey dots and diamonds show individual recruitments (we use two different symbols to highlight those values that are beyond one standard deviation); boxes cover the 2nd and 3rd quartile and whiskers extend 1.5 times the interquantile range (whiskers are asymmetric, comprising the most extreme observed values that are within 1.5 × IQR from the upper or lower quartiles). Parameters as in Fig. 11.

### 3.2.4 The Impact of the Input Current Strength on the Recruitment Time

Following the same approach used to obtain the results shown in Fig. 8 for a healthy subject, we present here an analysis on the impact of the stimulation strength on the recruitment mechanism. Fig. 17 displays the recruitment times of the first ten recruited areas using different amplitudes *I*_S_ of the step current *I*_S_(*t*), while fixing the duration *t*_I_ = 0.4 s. The analysis has been performed for patients CJ (panel A), CM (panel B), FB (panel C) and PG (panel D), thus integrating the information on the dependency on topological measures presented in the previous section. As expected, the recruitment times decrease for larger amplitudes. However, the order of recruitment does not substantially change. This implies that, whenever we increase the amplitude, the recruitment mechanism remains unaffected: the same populations are involved in the seizure spreading and in the same order. What changes is the speed of the spreading and the time necessary to observe a generalized seizure-like event, which is smaller for stronger currents. As a general remark, the brain areas that are recruited after the first ones (i.e. the 5*th*, 6*th*,…,10*th* recruited areas), tend to be recruited more simultaneously for increasing *I*_S_, thus leading to possible changes in the recruitment order. This can be appreciated especially for patient CJ: For an amplitude *I*_S_ = 10, for example, the 10*th* brain area (pink) gets recruited later than the 9*th* area (darkblue), while for a very strong currents (*I*_S_ = 100) the darkblue area gets recruited latest whereas the pink area gets recruited earlier.

**Figure 17.**
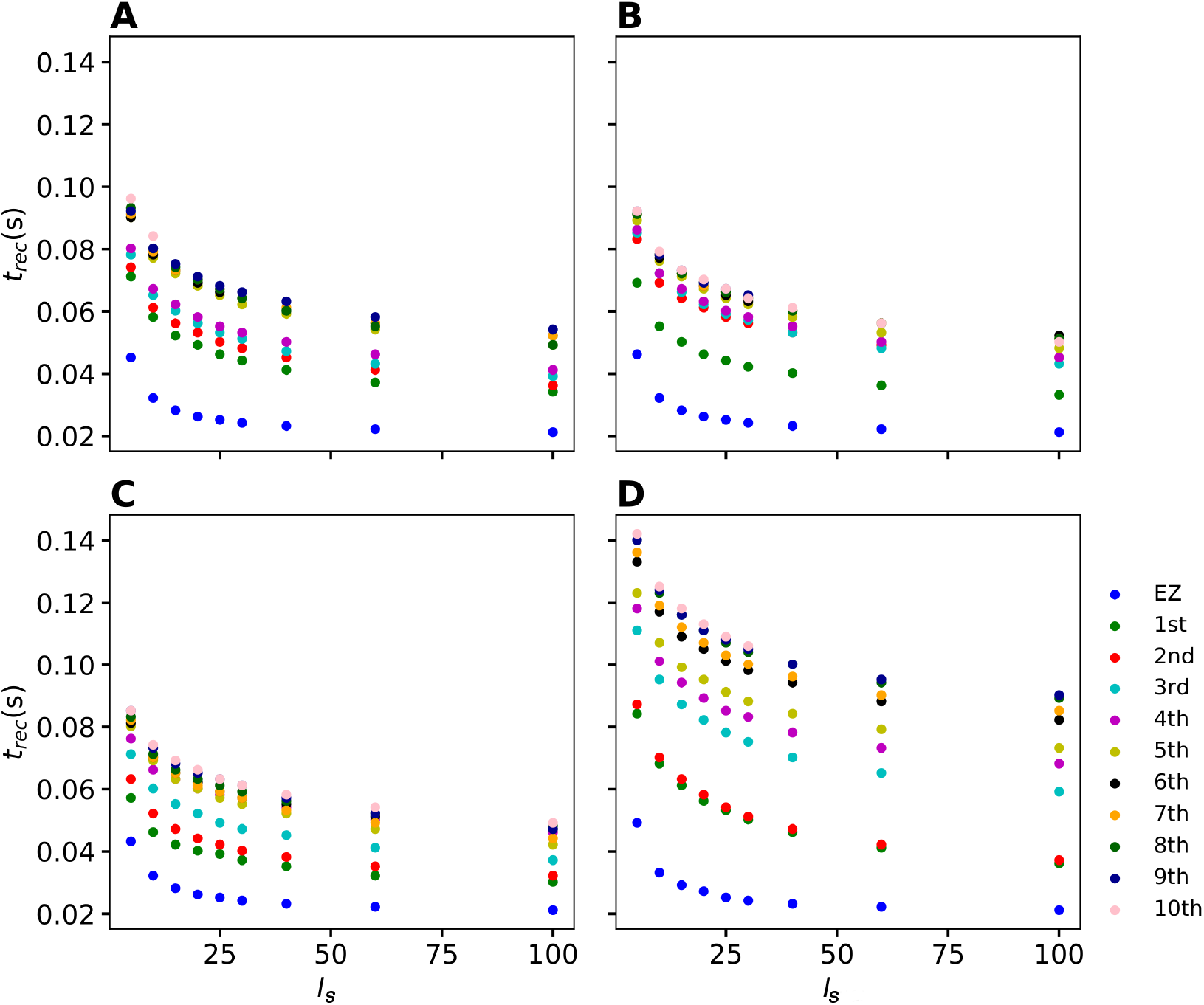
Recruitment times of the first 10 recruited areas as a function of the input current *I_S_* for the epileptic patients A) CJ, B) CM, C) FB and D) PG. The strength of the input current is varied between 0 and 100 on the x-axis while its duration is kept unchanged at *t_I_* = 0.4 s with respect to the previous numerical experiments. The order of the recruitment is color coded for each current strength (i.e. blue dots indicate the recruitment of the EZ, green dots indicate the first recruited area, red the second, etc.) and it holds the same for all investigated patients. Parameters as in Fig. 11.

On the other hand if we vary the step current duration *t*_I_ keeping the amplitude *I*_S_ = 15 fixed, we do not observe any change in the recruitment times of the first 10 recruited areas, analogously to the healthy subject case presented in Fig. 20 of the Supplementary Material. Results not shown.

**Figure 18.**
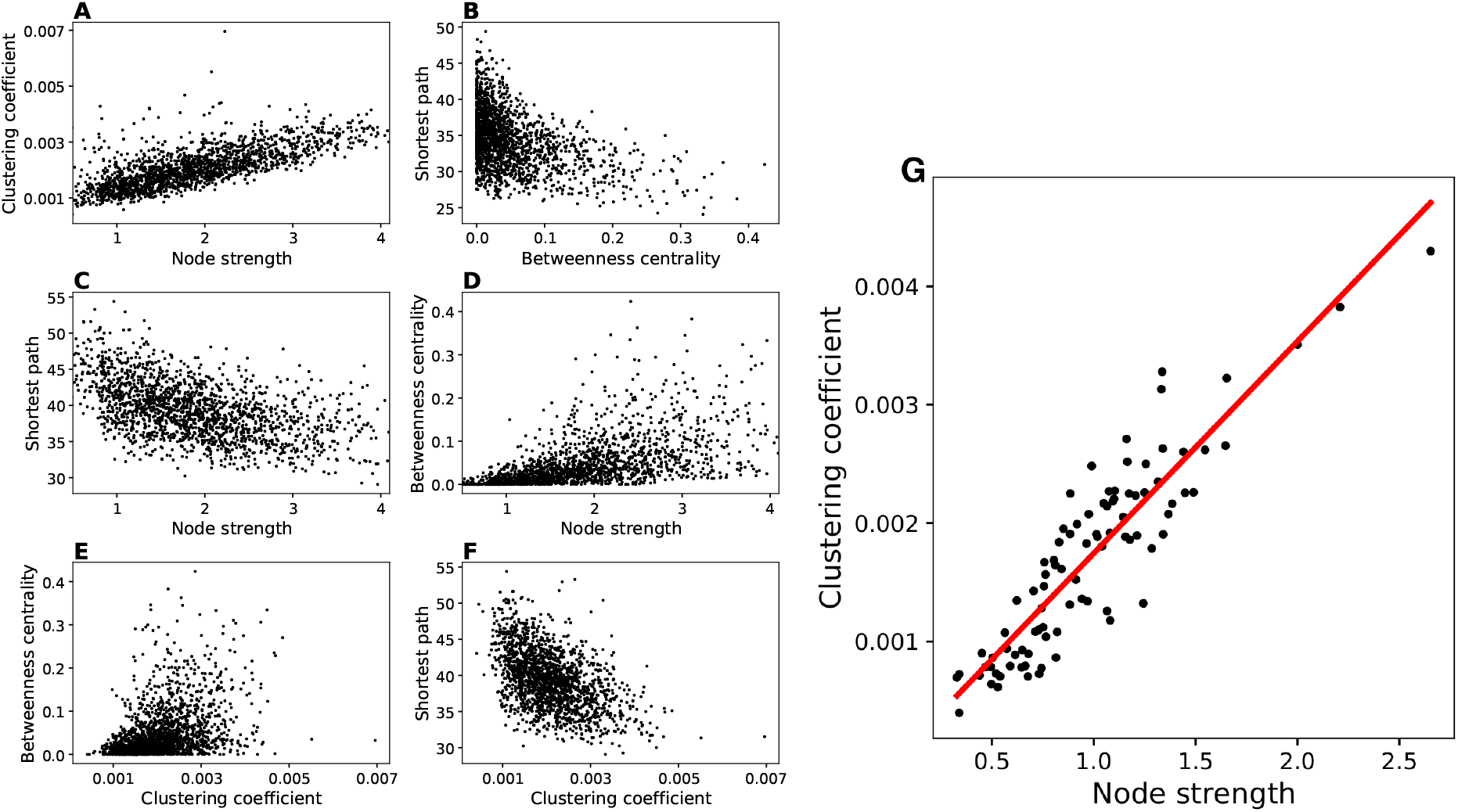
Network measure correlations of healthy subjects. Panels A-F are obtained plotting independently all node values for all the subjects (90*20=1800 data points). Panel G: Data are averaged over all 20 subjects. The single node values are averaged over the different subjects and afterwards, the correlation between node strength and clustering coefficient is estimated. Infinite values were excluded. The Pearson correlation of clustering coefficient and node strength of the averaged healthy DTI topology is *r* = 0.9 and much stronger compared to the average of all individual topologies *r* = 0.75 from panel A.

**Figure 19.**
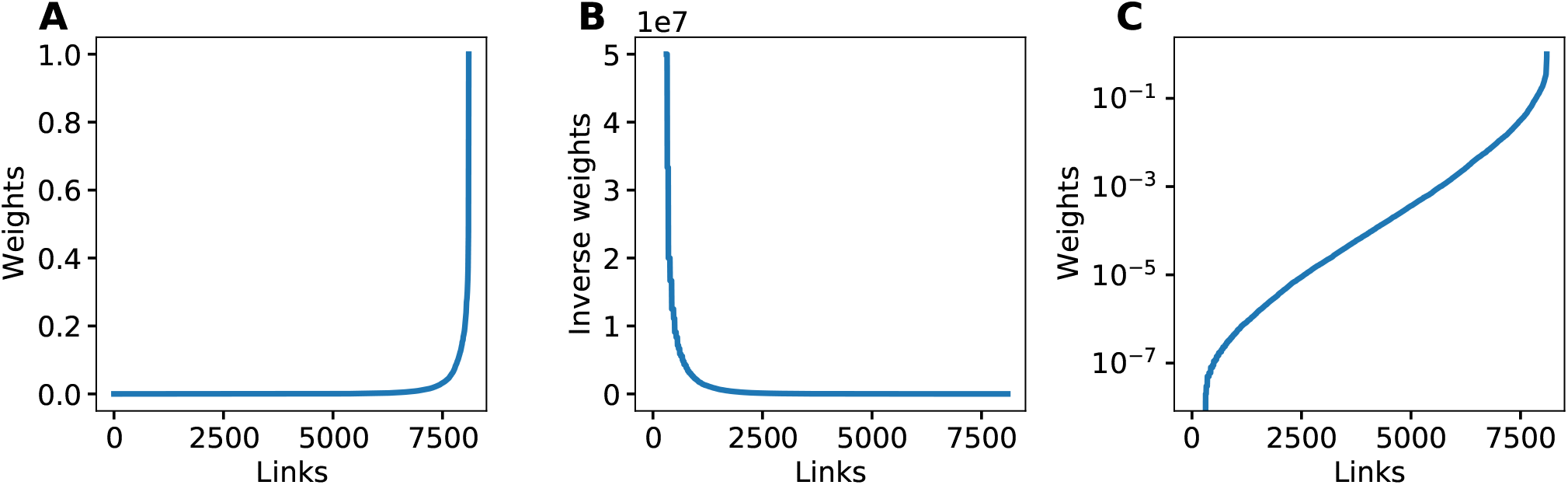
Weight Distribution of the DTI graphs. The weight distribution with weights on the x-axis in ascending order. a) Weight distribution, b) the inverse weight distribution, c) and the logarithmic weight distribution of the healthy averaged DTI graph. Note that c) matches the curve of recruitment times in 7 a).

**Figure 20.**
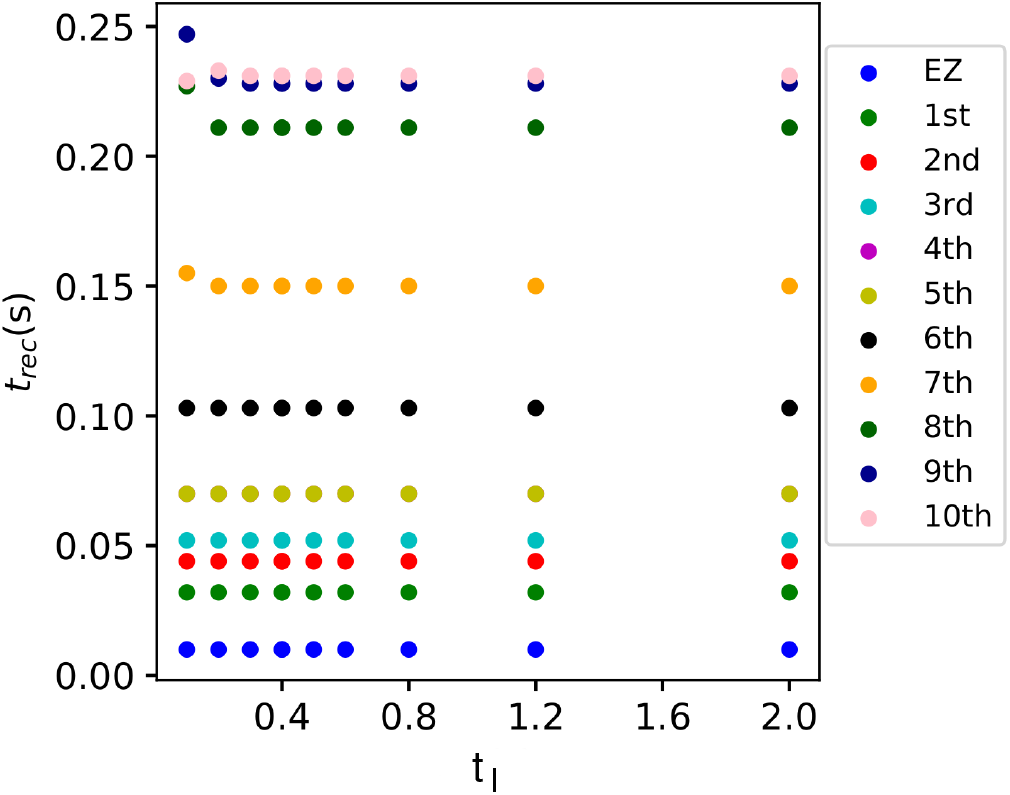
Input Current Duration Variation. Dependence of the recruitment time on the current duration *t_I_* while the current strength is kept constant at *I_S_* = 15. The y-axis shows the recruitment times of the first 10 recruited areas for each current strength. Blue is the EZ, green is the first recruited area, red the second, etc. The recruitment times are independent of the pulse duration. Parameters: *N* = 90, *σ* = 1, Δ = 1, 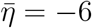, *I*_S_ = 15, stimulation site: brain area *k* = 45 for the healthy subject 0.

## 4 DISCUSSION

Neural mass models have been actively used since the 1970s to model the coarse grained activity of large populations of neurons and synapses Wilson and Cowan (1972); Zetterberg et al. (1978). They have proven especially useful in understanding brain rhythms Da Silva et al. (1974, 1976); Sotero et al. (2007), epileptic dynamics Wendling et al. (2016); Jirsa et al. (2014), brain resonance phenomena Spiegler et al. (2011), resting state Ghosh et al. (2008); Deco et al. (2011), task activity Huys et al. (2014); Kunze et al. (2016), neurological and psychiatric disorders Bhattacharya and Chowdhury (2015) and are very popular in the neuroimaging community Valdes-Sosa et al. (2009); Moran et al. (2013). Moreover, the desire to understand large scale brain dynamics as observed using EEG, MEG and fMRI has prompted the increasing use of computational models Bojak and Breakspear (2014). Large-scale simulators such as The Virtual Brain (TVB) Sanz-Leon et al. (2015) and research infrastructures such as EBRAINS (http://ebrains.eu) make heavy use of networks of interconnected neural mass models and enable non-expert users to gain access to expert state-of-the-art brain network simulation tools.

Although motivated by neurobiological considerations, neural mass models are phenomenological in nature, and cannot hope to recreate some of the rich repertoire of responses seen in real neuronal tissue. In particular their state variables track coarse grained measures of the population firing rate or synaptic activity. At best they are expected to provide appropriate levels of description for many thousands of near identical interconnected neurons with a preference to operate in synchrony, but they cannot reproduce the variation of synchrony within a neuronal population which is believed to underlie the decrease or increase of power seen in given EEG frequency bands. Importantly, unlike its phenomenological counterpart, the next generation neural mass model we have implemented in this paper, is an exact macroscopic description of an underlying microscopic spiking neurodynamics, and is a natural candidate for use in future large scale human brain simulations. The alternative method to heuristic neural mass models employed so far consists in performing large numerical simulations. Since the next generation neural mass model allows to overcome the limitations in the maximal affordable number of simulated neurons, it solves also the problems that are usually encountered in the analysis of spiking neural circuits addressed through numerical simulations, i.e. the limited available numerical resources.

In addition to this, the inability of a single neural mass model to support event-related desynchronisation/synchronisation Pfurtscheller and Da Silva (1999) or to capture the onset of synchronous oscillations in networks of inhibitory neurons Devalle et al. (2017), reminds us that these phenomenological models could be improved upon. While building more detailed biophysically realistic models of neurons would increase the computational complexity and the difficulties to interpret the behaviour of very high dimensional models in a meaningful way, the next generation neural mass models here applied, are very much in the original spirit of neural mass modelling, yet importantly they can be interpreted directly in terms of an underlying spiking model. This exact derivation is possible for networks of quadratic integrate-and-fire neurons, representing the normal form of Hodgkin’s class I excitable membranes Ermentrout and Kopell (1986), thanks to the analytic techniques developed for coupled phase oscillators Ott and Antonsen (2008). This new generation of neural mass models has been recently used to describe the emergence of collective oscillations in fully coupled networks Devalle et al. (2017); Laing (2017); Coombes and Byrne (2019); Dumont and Gutkin (2019) as well as in balanced sparse networks di Volo and Torcini (2018). Furthermore, it has been successfully employed to reveal the mechanisms at the basis of theta-nested gamma oscillations Segneri et al. (2020); Ceni et al. (2020) and the coexistence of slow and fast gamma oscillations Bi et al. (2020). Finally it has been recently applied to modelling electrical synapses Montbrió and Pazó (2020), working memory Taher et al. (2020) and brain resting state activity Rabuffo et al. (2020).

In this paper we have extended the single next generation neural mass model derived in Montbrió et al. (2015) to a network of interacting neural mass models, where the topology is determined by structural connectivity matrices of healthy and epilepsy-affected subjects. In this way we coped non only with the macroscopic dynamics self-emergent in the system due to the interactions among nodes, but also with the various differences related to the patient-specific analyses.

In absence of external forcing, the phase diagram of the system as a function of the mean external drive 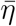 and synaptic weight *J* resembles this of the single neural mass model, since the same distinct regions can be observed: (1) a single stable node corresponding to a low-activity state, (2) a single stable focus (spiral) generally corresponding to a high-activity state, and (3) a region of bistability between low and high firing rate. However, when the system is subject to a transient external current, the scenario changes and is ruled by the interactions among different nodes. In this case, for low excitability values, a single stimulated node abandons the bistable region due to the applied current and it approaches, with damped oscillations, the high-activity state, which is a stable focus. On the other hand, for sufficiently high excitabilities, the single node stimulation leeds to the recruitment of other brain areas that reach, as the perturbed node, the high-activity regime by showing damped oscillations. This activity mimicks a seizure-like event and enables the modeling of propagation and recruitment: the seizure-like event originates in the EZ (as a results of the stimulation) and propagates to the PZ, identified by the other regions that fastly propagates the oscillatory activity. It is distinct from an actual seizure, which would require the emergence of self-sustained activity in the high-activity state Jirsa et al. (2014); Saggio et al. (2017, 2020)

The spectrogram analysis has revealed that the recruitment process is characterized by high frequency *γ* oscillations, thus reproducing the high-frequency (*γ*-band) EEG activity typical of electrophysiological patterns in focal seizures of human epilepsy. Many hypotheses have been formulated on the origin of this fast activity: (i) the behaviour of inhibitory interneurons in hippocampal or neocortical networks in the generation of gamma frequency oscillations Jefferys et al. (1996); Whittington et al. (2000); (ii) the nonuniform alteration of GABAergic inhibition in experimental epilepsy (reduced dendritic inhibition and increased somatic inhibition) Cossart et al. (2001); Wendling et al. (2002); (iii) the possible depression of GABA_*A,fast*_ circuit activity by GABA_*A,slow*_ inhibitory postsynaptic currents White et al. (2000); Banks et al. (2000); iv) the out of phase patterns of depolarizing GABAergic post-synaptic potentials onto pyramidal cells, generated by feed-forward activation of cortical interneurons Shamas et al. (2018). In any case high-frequency EEG waves originating from one or several brain regions are the most characteristic electrophysiological pattern in focal seizures of human epilepsy and can be observed, in our numerical experiments, both for healthy subjects and epileptic patients, though with a distinction: for the same excitability value, the activity takes place at higher frequency ranges in epileptic patients and it is mainly concentrated in the EZ. Even though it is not possible to exclude discrepancies partially imputable to the different scanning and preparation procedure of the structural connectivity matrices for the cohort of healthy and epilepsy-affected subjects, it turns out that the recruitment process is faster in epileptic patients, for which it is possible to observe generalize seizure-like events for smaller values of the excitability parameter 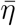. In particular, when comparing the results obtained for healthy subjects and epileptic patients, it turns out that the time necessary to recruit areas in the PZ is usually smaller for epileptic patients. However, the first recruited area is, in general, the area with the stronger connection to the EZ, independently of the considered structural connectivity matrix. The recruitment time in both cases is influenced by the strength of the external perturbation *I_S_*, and decreases for increasing strength, while no dependence is shown on the duration of the external perturbation.

More specifically for healthy subjects we have investigated the dependence of the recruitment mechanism on the single subject, in terms of the position of the eventual EZ and in terms of the topological measures of the single connectome. Brain network models of healthy subjects comprise 90 nodes equipped with region specific next generation neural mass models and each subject is characterized by a specific structural large-scale connectivity amongst brain areas. The smallest excitability values for which an asymptomatic seizure-like event occurs 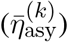 do not vary significantly from one subject to the other and do not show a relevant dependence on the stimulated area, while the smallest excitability values for which a generalized seizure-like event occurs 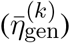, show fluctuations in the interval (−7, −5) for all stimulated nodes and for all the subjects. Nonetheless we have found many similarities at the level of topological measures, since there is always a strong correlation between 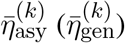 and node strength, clustering coefficient and shortest path, thus meaning that a region well connected is a region well recruited.

For epileptic patients, we have systematically simulated the individual seizure-like propagation patterns and validated the numerical predictions of the PZ against clinical diagnosis and SEEG signals. Patient-specific brain network models of epileptic patients comprise 88 nodes equipped with region specific next generation neural mass models and, for this set-up, we have studied the role of the large-scale connectome based on diffusion MRI, in predicting the recruitment of distant areas through seizure-like events originating from a focal epileptogenic network. We have demonstrated that simulations and analytical solutions approximating the large-scale brain network model behavior significantly predict the propagation zone as determined by SEEG recordings and clinical expertise, with performances comparable to previous analyses on this set of data Proix et al. (2017); Olmi et al. (2019), thus confirming the relevance of using a large-scale network modeling to predict seizure recruitment networks.

Most computational models of seizure propagation focus on small continuous spatial scales Hall and Kuhlmann (2013); Ursino and La Cara (2006); Kim et al. (2009) or population of neurons Miles et al. (1988); Golomb and Amitai (1997); Compte et al. (2003); Bazhenov et al. (2008); Chouzouris et al. (2018); Lopes et al. (2019); Gerster et al. (2020) while only small networks are commonly used to investigate the role of the topology and localization of the epileptogenic zone Terry et al. (2012). However functional, volumetric and electrographic data suggest a broad reorganization of the networks in epileptic patients Lieb et al. (1987, 1991); Cassidy and Gale (1998); Rosenberg et al. (2006); Bettus et al. (2009), thus laying the foundations for a different approach based on large-scale connectomes to identify the recruitment networks. The large-scale character of partial seizure propagation in the human brain has been only recently investigated, using patient-specific diffusion MRI data to systematically test the relevance of the large-scale network modeling in predicting seizure recruitment networks Proix et al. (2014, 2017, 2018); Olmi et al. (2019). In this framework of large-scale network modeling we can also place the results presented in this paper, since we have confirmed the importance of patient-specific connectomes to identify the recruitment process. As shown above, the topological characteristics of connection strength and shortest path play a non-trivial role in determining the spreading of seizure-like events, together with the localization of the epileptogenic zone, while the next generation neural mass model, here employed for the first time to study seizure spreading, allows us to construct patient-specific brain models via a multiscale approach: the variability of brain regions, as extracted from the human brain atlas, can be introduced in the mean-field parameters, thanks to the exact correspondence between microscopic and macroscopic scales guaranteed by the model itself. Improving the predictive power of the model by the means of anatomical data (available e.g. in the BigBrain and human brain atlas) will be the scope of further research.

## AUTHOR CONTRIBUTIONS

Moritz Gerster and Halgurd Taher performed the simulations and data analysis, writing original software and investigating the results. Data Curation is contributed by Viktor Jirsa, Maxime Guye, Fabrice Bartolomei, Antonín Škoch and Jaroslav Hlinka. All the authors validated the research and participated in the drafting process. Simona Olmi was responsible for conceptualization, supervision, state-of-the-art review (with Viktor Jirsa) and the paper write-up.

## DATA AVAILABILITY STATEMENT

All relevant data are within the paper and its Supporting Information files.

## SUPPLEMENTARY MATERIAL

**Figure 21.**
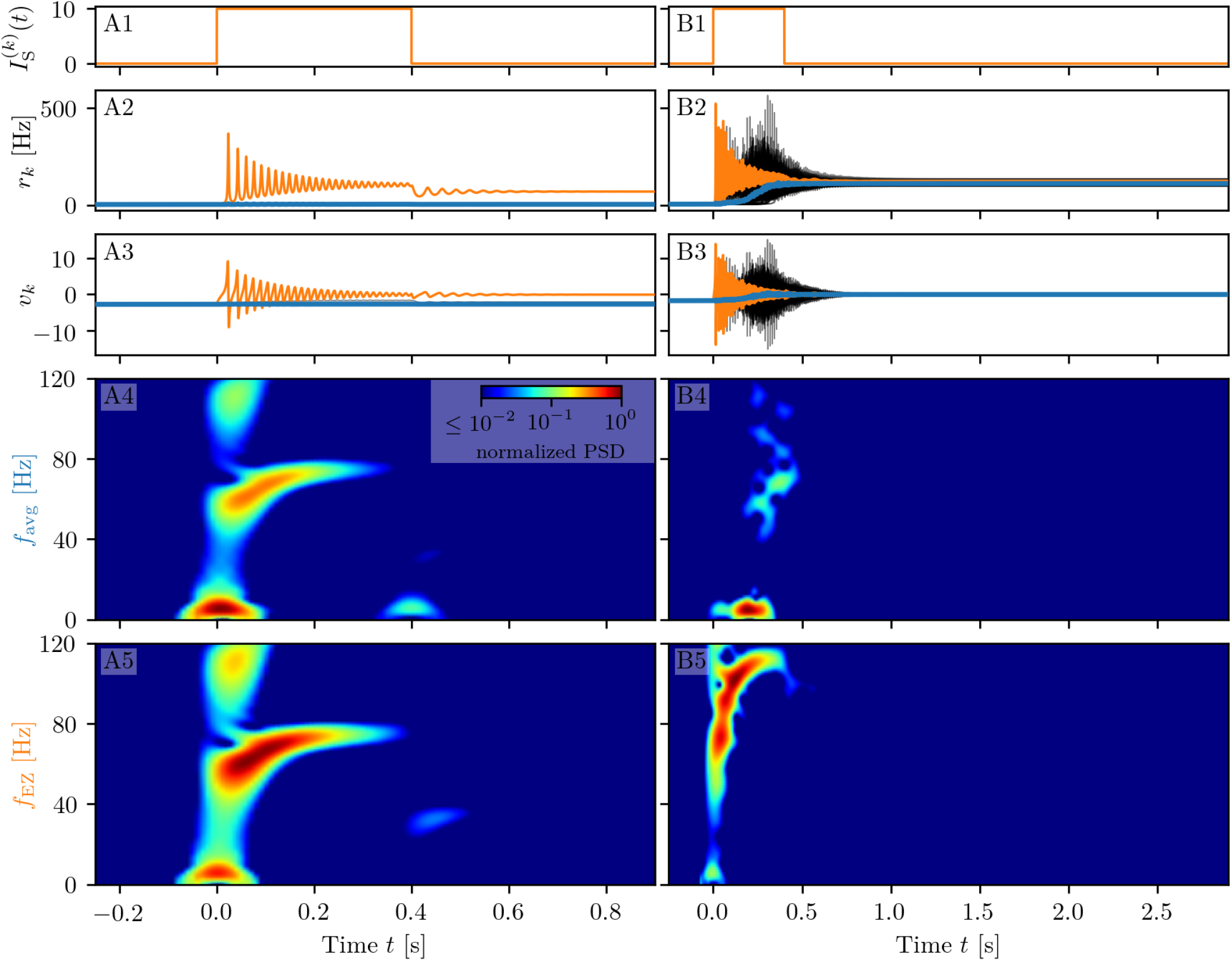
Spectrograms of mean membrane potentials for healthy subject sc2. (A1-B1) Stimulation current 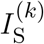, (A2-B2) population firing rates *r_k_* and (A3-B3) mean membrane potentials *v_k_* for the EZ (orange) and other populations (black). The blue curves show the network average firing rate and membrane potential. Non-stimulated node dynamics is plotted as transparent gray curves: some of the nodes adapt their voltage to the stimulation of the EZ and change during stimulation. However they do not reach the high-activity state regime. (A4-B4) Spectrogram of the network average membrane potential and (A5-B5) of the *v_k_* of the EZ. Column A shows an asymptomatic seizure-like event for 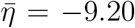, column B a generalized seizure-like event for 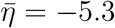. In both cases the EZ node 46 is stimulated. Parameter values: *N*_pop_ = 90, *τ*_m_ = 20 ms, Δ = 1, *J_kk_* = 20, *σ* = 1, 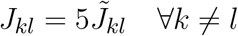.

**Figure 22.**
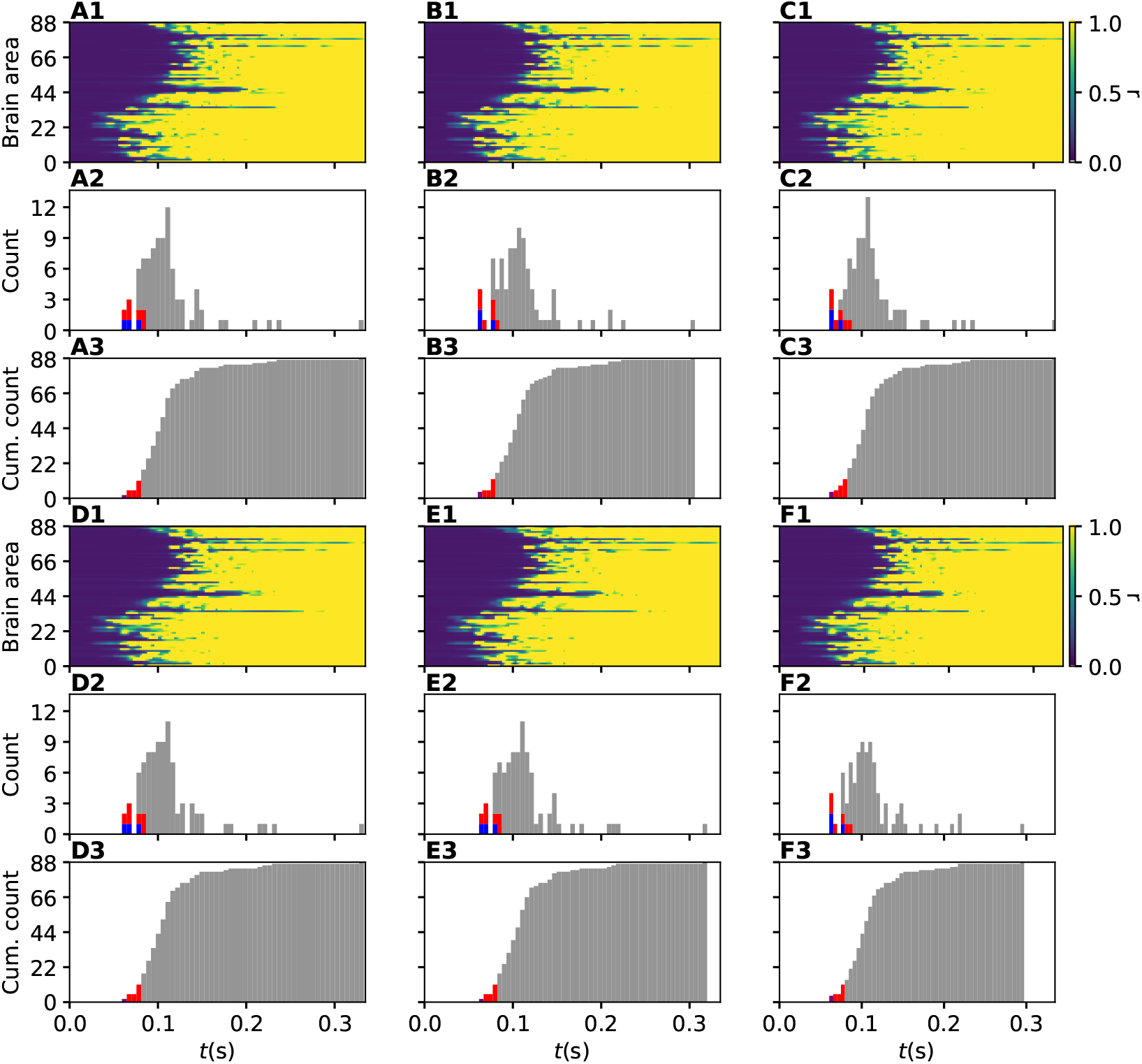
Recruitment times for patient CJ obtained for 6 different random Gaussian distributions of 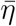. (A1-F1) Spacetime plots of the average firing rates of all brain areas. (A2-F2) Histograms of the recruitment times. Red (blue) bins identify those recruited area that belong to PZ_Clin_ (PZ_SEEG_). (A3-F3) Cumulative histograms of the recruitment times. Purple bin: EZ. Red bins: first 10 recruited areas. Parameters as in Fig. 11. For one exemplary patient, CJ, we show here in detail the impact of different realizations of 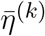, drawn from a Gaussian distribution (centred at 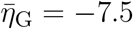 with standard deviation 0.1), on the recruitment times of the brain areas. In particular we have considered it to be sufficient to present results for six out of ten realizations, due to the large similarities between the outcomes. Space-time plots of the average firing rates give an immediate visualization of the recruitment events for each brain area. We find that the pattern of recruitment does not change substantially for different realizations of the 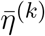. The EZ is localized in the area lh-LOCC, that corresponds to node *k* = 20: The firing rate of this population increases immediately upon stimulation, thus giving rise to the recruitment mechanism. The brain areas in the PZ are rapidly recruited: In general the first ten areas are always recruited in less then 0.1 s, followed by a continuous increase of the number of recruited nodes. Finally, it is worth noticing that the first recruited areas correspond to those predicted clinically.

**Figure 23.**
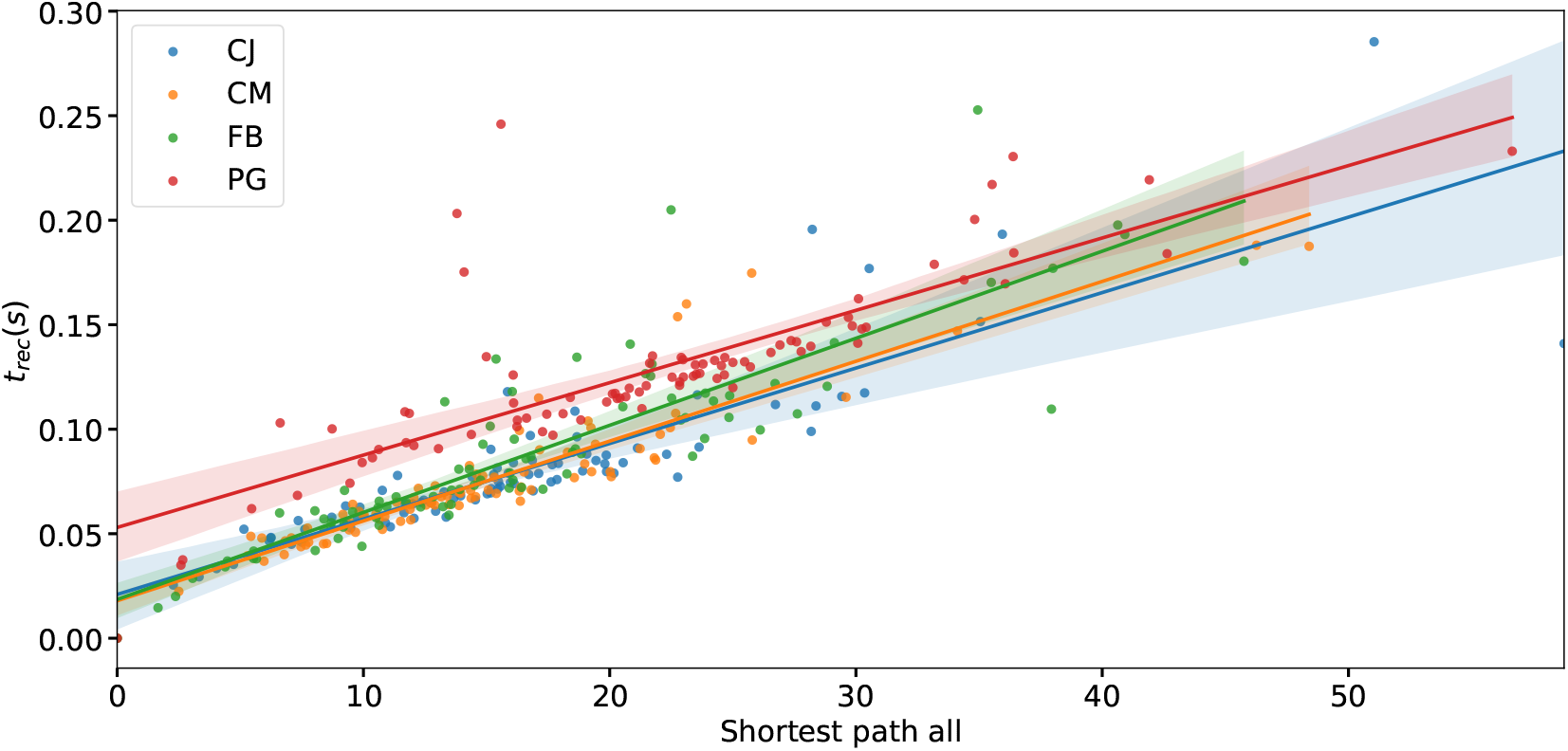
Recruitment time and Shortest Path. The recruitment times *t_rec_* as a function of the shortest path to the EZ are shown for four patients and all brain areas. Same af Fig. 14 A, with a regression fit that underlines the approximately linear relationship between the shortest path length and the recruitment time. Parameters as in Fig. 14.

## Footnotes

1 While the actual role of the specific regions might in reality be affected by other factors, not captured by the used structural connectivity estimate and the details of the current model, this highlights the effect of network structure on propensity to seizure-like events. The (para)hippocampal region is, in fact, one of the most commonly affected by epilepsy.

2 Please note that, irrespectively of the numerical results, any difference observed between the structural connectivity matrices obtained from the cohort of healthy subjects and epileptic patients may be (at least partially) ascribed to the different acquisition and processing procedures in the two research centers rather than due to disease-related causes.

